# Mechanisms shaping the transcriptome of E. coli to non-lethal rifampicin stress

**DOI:** 10.1101/2025.09.02.673683

**Authors:** M. M. Azevedo, A. M. Arsh, R. Jagadeesan, Andre S. Ribeiro

## Abstract

Rifampicin, by hampering transcription, perturbs bacteria even at non-lethal concentrations. In response, Escherichia coli adapts its phenotype to minimize mortality, which is followed by beneficial mutations. Most genome-wide transcriptional regulatory mechanisms controlling the adaptations remain unidentified. We studied the genome-wide, time-resolved, transcriptional program of susceptible E. coli cells under non-lethal rifampicin stress. Dynamically, the transcriptome widely diverged from the control, but later partially realigned. The mechanisms were changes in RNAP and Gyrase levels, promoter sequences, transcription factor network, intergenic distance, sensitivity to DNA supercoiling buildup, σ factor specificity, (p)ppGpp, and a few global regulators. These results show that the genome-wide response dynamics to rifampicin is influenced by the structure of the gene regulatory network. Next, we compared the evolutionarily distant pathogen Mycobacterium tuberculosis. In both species, adjacent genes on the DNA exhibited similar response strengths. Also, the response strengths of orthologous genes were correlated, suggesting that both species implement similar (likely beneficial) phenotypic adaptations. In support, E. coli orthologs were enriched in the mechanisms identified as influential. Overall, E. coli, and likely other bacteria, have mechanisms influencing specific gene cohort responses to non-lethal rifampicin stress, which likely enhances survivability, thus facilitating the emergence of resistance.

## 1. INTRODUCTION

In natural environments, exposure to non-lethal antibiotic (AB) stress is common. E.g., it occurs when the bacterium is distant from the AB source, causing the AB concentration to be lower than the minimum inhibitory concentration (1). Interestingly, these events can lead to the emergence of AB resistance mechanisms (2–7). Similarly, a few bacteria can trigger AB persistence, which consists of becoming physiologically dormant. Global regulators such as (p)ppGpp play a central role but several other molecular mechanisms can also be involved (8).

Since most ABs are toxic even at non-lethal levels, bacteria under non-lethal antibiotic (AB) stress can also trigger adaptation mechanisms, such as stress responses and metabolic changes to increase cell survivability (9–14). Escherichia coli, for example, can tune several cellular processes to increase survivability to non-lethal AB stress, as well as to enable survival to transient exposures to lethal AB concentrations, even when lacking resistance mechanisms (15, 16). The processes and mechanisms tuned include energy metabolism, drug efflux pumps (e.g. AcrAB-TolC), quorum sensing, (p)ppGpp signaling, and toxin-antitoxin systems (17–20), among other. For example, the tuning of iron acquisition, DNA repair, aerobic respiration, and carbon metabolism reduces the killing potency of rifampicin (21).

The gene regulatory mechanisms of E. coli responsible for triggering these advantageous phenotypic modifications remain unknown, potentially due to difficulties in tracking information propagation in the GRN. E.g., (22) reported an inconsistency between transcription factor (TF) input and output genes. Recently, as more regulatory mechanisms are dissected (23), it was possible to find a correlation between TFs and the dynamics of the GRN, under some genome-wide stresses (13, 24).

Rifampicin is an AB binds to RNAP holoenzymes, before (when free-floating) and/or after the RNAP binds to DNA and forms an open complex (but not after subsequent steps) (25, 26). Once bound, it prevents the RNAP from escaping promoters (25, 27), directly and quickly reducing RNA and corresponding protein levels globally, as they are mostly determined by transcription initiation kinetics (28). What happens after this initial genome-wide decrease when under non-lethal AB stress is yet to decipher, since studies of the genome-wide effects of non-lethal AB concentrations are uncommon. A recent study has provided evidence that non-lethal rifampicin concentrations cause significant signal propagations along operons (13).

We investigated which gene regulatory mechanisms influence the dynamics of the GRN of E. coli when under non-lethal concentrations of rifampicin. We exposed cells to rifampicin and performed time-resolved RNA-seqs to, first, identify responsive gene cohorts. Next, we studied which regulatory mechanisms influenced the genes’ behavior over time. Namely, we assessed how RNAP, promoter sequences, transcription factors, activity of neighboring genes, DNA supercoiling buildup, σ factors, (p)ppGpp, and global regulators of transcription shape the genes response timing and magnitude.

Finally, we studied Mycobacterium tuberculosis genome-wide response to non-lethal rifampicin stress, with emphasis on orthologous genes. This bacterium is estimated to be carried by almost 2 billion people in 2015 (29) while Rifampicin is one of the most widely used ABs to treat it, in combination therapy (30). Dissecting the features that could enhance their survival to rifampicin could contribute to global health and evaluate the extent to which the results for E. coli are generalizable to other bacterial species.

## 2. METHODS

### 2.1 Bacterial strains

MG1655 E. coli cells were used as the wild type (WT) strain to collect RNA-seq and other data, unless stated otherwise. A YFP fusion strain library (31) was used to measure several single-cell protein levels, including GyrA, GyrB, spoT, and RpoB. An RL1314 strain with RpoC endogenously tagged with GFP (generously provided by Robert Landick, of the University of Wisconsin-Madison) was used to measure RpoC levels. Finally, the global regulators MarA, Fnr, LexA, Fis, and SoxS were measured using a strain library of single-copy plasmids with the promoters of interest controlling the expression of GFP (32).

Finally, a strain library generously provided by the Elf lab (33, 34) was used to assess effects from changes in promoter sequences. The strains have a chromosomal integrated pFAB120 synthetic promoter, followed by one LacI binding site that differs by one to a few nucleotides from the same binding site in the other strains. This affects the association-dissociation kinetics of the repressor, LacI (34). The construct starts with a promoter (35 bp long) immediately followed by one operator (20-21 bp long), a double ribosome binding site, RBS (33 bp long), a mVenus coding sequence (717 bp) and a double terminator sequence (57 bp). The mutations are located at the operator, +3 to +21 bp downstream of the TSS. Thus, they are within +1 to + 60 downstream from the TSS. Arguably, they may interfere with RNAP-promoter escape, but not with RNAP binding to the TSS.

Importantly, the mutations, being within +1 to +20 nucleotides downstream from the TSS, might affect RNAP escape from the TSS, which is the step blocked by rifampicin (25). Contrarily, they should not affect RNAP binding affinity (34). Due to this, if the promoters are fully induced (> 500 µM IPTG), the mutations should not be influential since full induction implies that LacI’s rarely (or never) bind to the binding sites. Similarly, if the promoters were fully repressed (0 µM IPTG), the mutations might also not be influential, since too few RNAs could be produced (unless the mutations caused a strong weaking in binding affinity). As such, to detect differences with the mutations, we compared the promoters’ response with and without rifampicin, while half-induced with 250 µM, by flow-cytometry (Methods section 4.5).

We used 7 out of the 35 synthetic promoters. For simplicity, we selected a set that changed linearly in the ratio between LacI’s association and dissociation rate to its regulatory site as reported in (34). This should facilitate interpreting the results of adding rifampicin.

When subjecting cells to rifampicin, we measured protein levels 50 minutes after adding both IPTG (half-induction concentrations) and rifampicin. Then, we again measured proteins, at 90 minutes after adding IPTG. This timing was selected as follows. The first 50 minutes should suffice for IPTG to enter cells and activate promoters (∼10 minutes) (35), as well as for increasing the corresponding protein levels, given the fast rates of RNA production (∼10 min-1) and protein production and maturation (mVenusNB, 4.7 min-1 at ∼30°C (36)). Meanwhile, the subsequent 40 minutes should suffice for protein levels to become lower than the control, given the intake time of rifampicin (5 min) and degradation rate (0.8 min−1) of the RNAs coding for the proteins (34). For control experiments, for each strain, we performed the same measurements without adding rifampicin.

### 2.2 Growth conditions and rifampicin

In general, from glycerol stocks at -80 °C, we streaked cells on lysogeny broth (LB) agar plates with ABs and kept at 37 °C overnight. Next, a single colony was picked, inoculated into fresh LB medium and kept at 37 °C overnight with appropriate ABs and aeration at 250 rpm. From overnight cultures, we diluted cells (1:1000 in LB media) and then incubated them at 37 °C with aeration. Finally, cells grew until reaching an optical density of ∼0.3 at 600 nm (OD_600_) with rifampicin being added at that moment. IPTG, when added, was when cells reached ∼0.3 OD_600_, then 50 minutes passed, at which point we added rifampicin. Measurements were conducted 40 minutes after that.

### 2.3 Minimum inhibitory concentration

We measured minimum inhibitory concentration as in (15). Overnight cultures were diluted 1 in 10000 in fresh LB media, grown with 2-fold dilutions of rifampicin and incubated for 24 h at 37 °C and 250 rpm shaking. We measured OD_600_ with a Bioteks Synergy HTX Multi-Mode Reader.

### 2.4 RNA-seq

We performed RNA-seqs at 80 and at 160 minutes after adding rifampicin, in addition to the control. The data for 160 minutes has been used in (13), while the (critical) data for 80 minutes is first published here. The RNA-seq of early responses to Novobiocin were performed 120 minutes after adding Novobiocin. These additional 40 minutes are in line with the slower permeation rates of Novobiocin, compared to rifampicin (37), through *E. coli* cell walls. Further, the effects of Novobiocin will only be detected after DNA supercoiling is affected, while rifampicin acts directly on transcription (14). The RNA-seq data related to Novobiocin was published in (14). Both measurements and data analysis were performed using the same methodology as in (13) described next.

Sample preparation: cells in the mid-exponential growth phase (OD_600_ ∼0.3) were subject to rifampicin (2.5 ng/μL). At 80 and 160 minutes later, respectively, we collected cells from 3 independent colonies. We also collected cells from colonies not subject to AB (control). After that, 5 mL of the culture was treated with a double volume (10 mL) of RNA protect bacteria reagent (Qiagen, Germany) for 5 minutes at room temperature, to prevent RNA degradation. Next, cells were pelleted and frozen at -80 °C. The next morning, total RNA was extracted using the RNeasy kit (Qiagen, Germany).

Sequencing: The RNA was treated twice with DNase (Turbo DNA-free kit, Ambion, USA) and quantified using Qubit 2.0 Fluorometer RNA assay (Invitrogen, Carlsbad, CA, USA). The total RNA quality was determined by 1% agarose gel staining with SYBR Safe (Invitrogen, Carlsbad, CA, USA), using UV in a Chemidoc XRS imager (Biorad, USA). RNA integrity was measured by Agilent 4200 TapeStation (Agilent Technologies, Palo Alto, CA, USA). Finally, RNA library preparations, sequencing, and quality control of sequenced data was performed by GENEWIZ, Inc. (Leipzig, Germany). Meanwhile, sequencing libraries were multiplexed and clustered on one lane of a Flowcell, which was loaded on an Illumina NovaSeq 6000 instrument. The samples were sequenced using a single-index 2×150 Paired-End (PE) configuration. Image analysis and base calling were conducted by the NovaSeq Control Software v1.7 (Illumina NovaSeq). Raw sequence data (.bcl files) was converted to “fastq” files and de-multiplexed using Illumina bsl2fastq v.2.20. One mismatch was allowed for index sequence identification.

Data analysis pipeline: Step 1: RNA sequencing reads were trimmed to remove adapter sequences and nucleotides with poor quality using Trimmomatic v.0.39 (38); Step 2: To generate BAM files, trimmed reads were mapped to the reference genome E. coli MG1655 (NC_000913.3) using Bowtie2 aligner v.2.3.5.1 (39); Step 3: Unique gene hit counts were calculated using ‘featureCounts’ from the ‘Rsubread’ R package (v.2.8.2) (40). Genes with less than 5 counts in more than 3 samples, and genes whose mean counts were smaller than 10 were removed; Step 4: The DESeq2 R package (v.1.34.0) (41) was used to calculate log2 fold changes (LFC) of RNA read counts between conditions and corresponding p-values, using Wald tests (function ‘nbinomWaldTest’).

Because some genes were removed from the analysis, some gene cohorts differ in size between 80 and 160 minutes.

### 2.5 Flow cytometry

We used an ACEA NovoCyte Flow Cytometer and Novo Express software (ACEA Biosciences Inc.). Cells are diluted (1:10000) in 1 mL of PBS solution and vortexed for 10 seconds. We collect 3 biological replicates per condition, 50000 cells per replicate. Flow rate is set to 12 µL/minute.

To detect GFP, YFP, and mVenusNB, we use the FITC channel (-H parameter) with 488 nm excitation, 530/30 nm emission, and 14 μl/min sample flow rate with a core diameter of 7.7 μm. PMT voltage was set to 600 for FITC, and sample flow rate to 14 μl/min, with a core diameter of 7.7 μm. PMT voltage was 584. To remove data from particles smaller than bacteria, the detection threshold was set to 5000. We also collected the ‘pulse width’ to use as a proxy for cell size (42, 43). No gating was applied. All events were collected by the Novo Express software from ACEA Biosciences Inc.

### 2.6 Spectrophotometry

We monitored cell growth from optical density at 600 nm with a Biotek Synergy HTX Multi-Mode Reader.

### 2.7 Cellular ATP

We grew QUEEN-2m cells, a kind gift from Hiromi Imamura of Yamaguchi University in Japan (44), in standard growth conditions, with and without rifampicin. We tracked the cells’ ATP levels using a Biotek Synergy HTX Multi-Mode Reader, exciting the solution at 400 nm and recording the emission at 513 nm. Then, the solution was re-excited at 494 nm and the emission was recorded at 513 nm. The ratio between the 513 nm emission intensities at the two excitation wavelengths, denoted ‘400ex/494ex’, is used as a proxy for cell ATP levels (44).

### 2.8 AT richness and p distance

When quantifying AT richness, we started by considering all promoters identified in RegulonDB (23). We then discarded 37% of them, since their DNA sequence was not available. Of the remaining 1057 promoters, we considered their DNA sequence from 60 nucleotides upstream of the TSS up to 20 nucleotides downstream of the TSS. Assuming the TSS in position +1, from the nucleotides in positions -60 to +20 (12, 75), we calculated the fractions of A, C, G and T, respectively. The promoter AT-richness was estimated by summing the fractions of A and T in this region (45).

Meanwhile, the p-distance of a promoter (46) is the fraction of its nucleotides between positions −41 to −1 (with the TSS starting at +1) that differ from the consensus (most common) nucleotide in that position of a gene cohort. Since measurements were performed with cells in the exponential growth condition, we considered the cohort of genes whose promoters have preference for σ^70^. The same 1057 promoters were considered as for the AT richness.

### 2.9 Statistical tests

p-values were obtained from 2-sample T-tests to assess if two distributions are from independent random samples from normal distributions with equal means and equal, but unknown, variances. For p-value < 0.05, we rejected the null hypothesis that the distributions have equal means.

## 3. RESULTS

### 3.1 Timing of the main GRN-dependent events triggered by rifampicin

Due to its capacity to mechanically block transcription, rifampicin has been used to measure global RNA degradation rates by microarray (47–52) and RNA-seq (28, 53–55). To be precise, RNA production needs to be entirely halted. That requires lethal rifampicin concentrations (50 – 500 µg/mL). Thus, it is not possible to study information propagation in the GRN. Meanwhile, in (56), cells were allowed to gain resistance and, thus, survived. However, rifampicin no longer affected the GRN as in AB susceptible cells, since the resistance is gained by random point mutations that hamper rifampicin’s binding affinity to RNAP (25, 57).

Figure 1 illustrates the timeline of the events expected to be triggered by non-lethal rifampicin concentration, given its mode of action (MoA) (25) and the effects of reducing the free-floating RNAP concentration (24). First, rifampicin should enter cells quickly after added to the medium (less than 5 minutes (58)). Its MoA is binding to RNAP and blocking promoter escape, hampering RNA production (25). Thus, the RNA numbers of active genes should decrease. In E. coli, ∼50% of the genes should be active during exponential growth (31).

**Figure 1:**
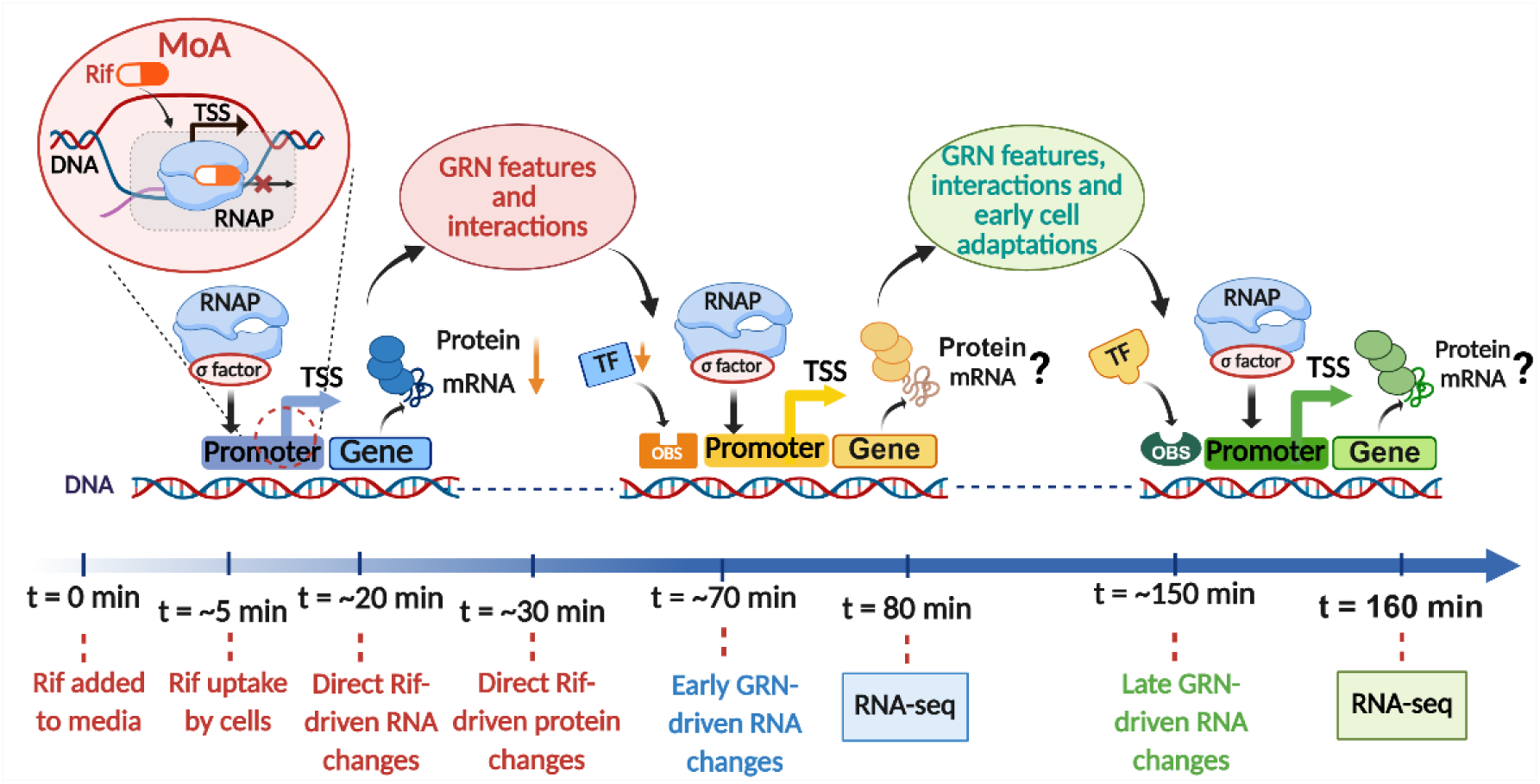
Illustration of the expected timeline of events on the GRN of *E. coli* triggered by rifampicin. Image adapted from (24). We predict three main phases: (i) First, genome wide changes in the transcriptome (reductions in RNA and TF levels), directly caused by rifampicin (due to blocking transcription initiation of active genes); (ii) A second, early GRN-driven phase of genome-wide changes in the transcriptome, largely already driven by GRN features and interactions (such as TF-operator binding site (OBS) interactions). Finally, (iii) a late GRN-driven phase influenced by GRN features, as well as early phenotypic adaptations, implemented during the previous phase (e.g. increased efflux pumps numbers (62)). Is it noted that this illustration only provides a simplified perspective of the flow of information, since many complex contributions (e.g. TF interaction strengths, promoter strengths, global regulators, (p)ppGpp, DNA supercoiling, etc.) are all represented under “GRN features”.

To estimate how long it takes for the RNA numbers to first change, at a genome-wide level, we considered that cells have only 10 or less RNAs per active gene in the exponential growth phase (31). Meanwhile, RNA half-lives are, on average, ∼6 min long (31, 47). Thus, RNAs can quickly become (nearly) absent in ∼15 min.

Next, we considered corresponding protein decay rates will be affected by degradation and dilution by cell division (59). If cells are in the exponential growth phase, we estimated that protein numbers could change within ∼30-40 minutes after adding rifampicin (31, 36, 60, 61).

Subsequently, these initial changes of the transcriptome and proteome, largely reductions, should be followed by a phase of global transcriptome changes, positive and negative, that are largely driven by the GRN (24). We named it ‘early GRN-driven phase’. We expected that the transcription factor network (TFN) will be particularly influential (Figure 1). Other GRN features (e.g. DNA supercoiling and global regulators) should also be influential (13, 24).

Given the expected times for RNA and protein numbers to change following the original changes, to study how the GRN modifies the initial genome-wide effects, we performed an RNA-seq at 80 min after adding rifampicin (Figure 1).

Finally, since we applied rifampicin below the minimum inhibitory concentration (MIC), *E. coli*’s *basal* AB adaptation mechanisms (e.g. increases in efflux pumps (62) and RNAP (63) levels) might suffice to reduce the AB effects over time. Namely, these natural mechanisms might contribute to a third ‘Late GRN-driven phase’ of transcriptomic changes where there is a partial return to pre-AB states. To also study this phase, we performed an additional RNA-seq, at 160 minutes after adding rifampicin (Figure 1).

### 3.2 Rifampicin effects on cell growth rate, size, ATP and, RNAP

To study how the GRN affects the propagation of the initial effects of rifampicin, we selected the maximum rifampicin concentration that *does not* affect cell growth or morphology. Growing cells as described in Methods Section 4.2, the maximum rifampicin concentration not influencing the growth rate significantly was 2.5 µg/mL (Figure 2A), in agreement with (63).

**Figure 2.**
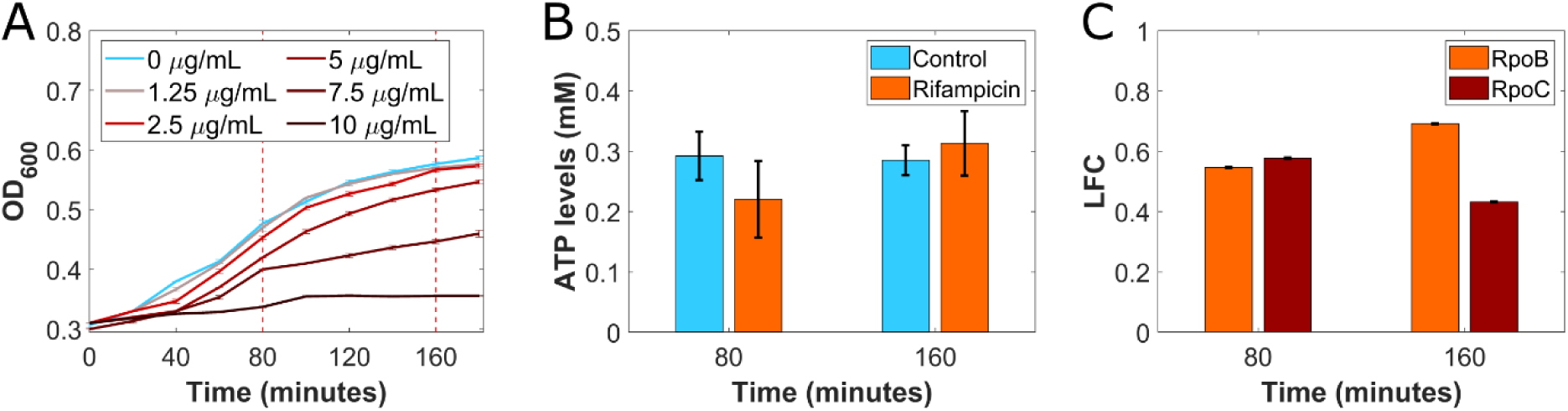
Time-lapse effects of rifampicin. **(A)** Cell growth as measured by OD_600_ after adding various rifampicin concentrations, respectively. Error bars calculated with the standard error of the mean (SEM) from 3 biological replicates. **(B)** ATP levels (Methods Section ‘Cellular ATP levels’) at 80 and 160 minutes in cells under 2.5 µg/mL of rifampicin and in control conditions as measured by spectrophotometry. Error bars calculated using the standard error of the mean (SEM) from 3 biological replicates and the propagation of uncertainty method. **(C)** Mean single-cell expression levels of two subunits of RNAP (RpoB-YFP and RpoC-GFP, respectively) in cells under 2.5 µg/mL of rifampicin relative to control conditions, as measured by flow-cytometry. LFC stands for Log_2_(fold change relative to the control condition). Error bars calculated using the standard error of the mean (SEM) from 3 biological replicates and the propagation of uncertainty method.

For comparison, the MIC (Methods Section 4.3) is 7.5 µg/mL (3× higher), similar to previous reports (64) (12.5 µg/mL or lower).

We also tracked ATP (Methods section 4.7) by spectrophotometry (Methods Section 4.6), as its depletion could damage the GRN structure and functioning. Specifically, lack of ATP can hamper the energy-consuming process of DNA supercoiling regulation. That, in turn, can hamper transcription and DNA replication (65). However, 2.5 µg/mL of rifampicin did not change ATP levels significantly (Figure 2B).

We also estimated changes in cell size using the flow-cytometry parameter ‘pulse width’ as a proxy (43) (Methods section 4.5). The size relative to the control only increased by 4% in 120 minutes (p-value of a 2-sample t test < 0.05 (Methods section 4.9)).

Finally, non-lethal rifampicin levels have been reported to increase transiently the amount of RNAP β and β’ subunits in *E. coli* (66–68) and in *M. tuberculosis* (69). To confirm this, we measured RpoC:GFP in RL1314 cells under rifampicin (Methods section 4.1). RpoC levels were significantly higher than the control (Log_2_(fold change), LFC > 0) at both 80 and 160 minutes, as measured by flow-cytometry (Figure 2C).

We validated this finding using another strain, carrying RpoB-YFP (Figure 2C) (Methods section 4.1). Visibly, RpoB levels are higher than the control (LFC > 0). These conclusions are further supported by RNA-seq data, shown below. The increased RpoB and RpoC levels could explain, at least partially, some of the subsequent changes in RNA levels reported below.

### 3.3 Non-lethal rifampicin stress affects the transcriptome transiently, causing diverse single-gene responses

Approximately ∼52% of all TF-gene interactions of *E. coli* have been identified as activations and ∼47% as repressions (23, 24). Thus, the TFN might convert the initial genome-wide reductions in RNA numbers into similar numbers of activations and repressions, at least in genes regulated by TFs. This ‘signal propagation’ is also influenced by many other variables such as global regulators, etc., as well as by each TF interaction strength, making it highly heterogeneous.

We analyzed the transcriptome at 80 and 160 minutes after adding rifampicin (2.5 µg/mL) (Methods Section 4.4). Figure 3A shows the distributions of the log₂ fold change (LFC) of all genes, relative to the control at the same time point. At 80 minutes, the log₂ fold change (LFC) distribution is broad, with many genes showing strong responses. In contrast, at 160 minutes, the GRN appears to ‘recover’, i.e., differing less from control cells.

**Figure 3.**
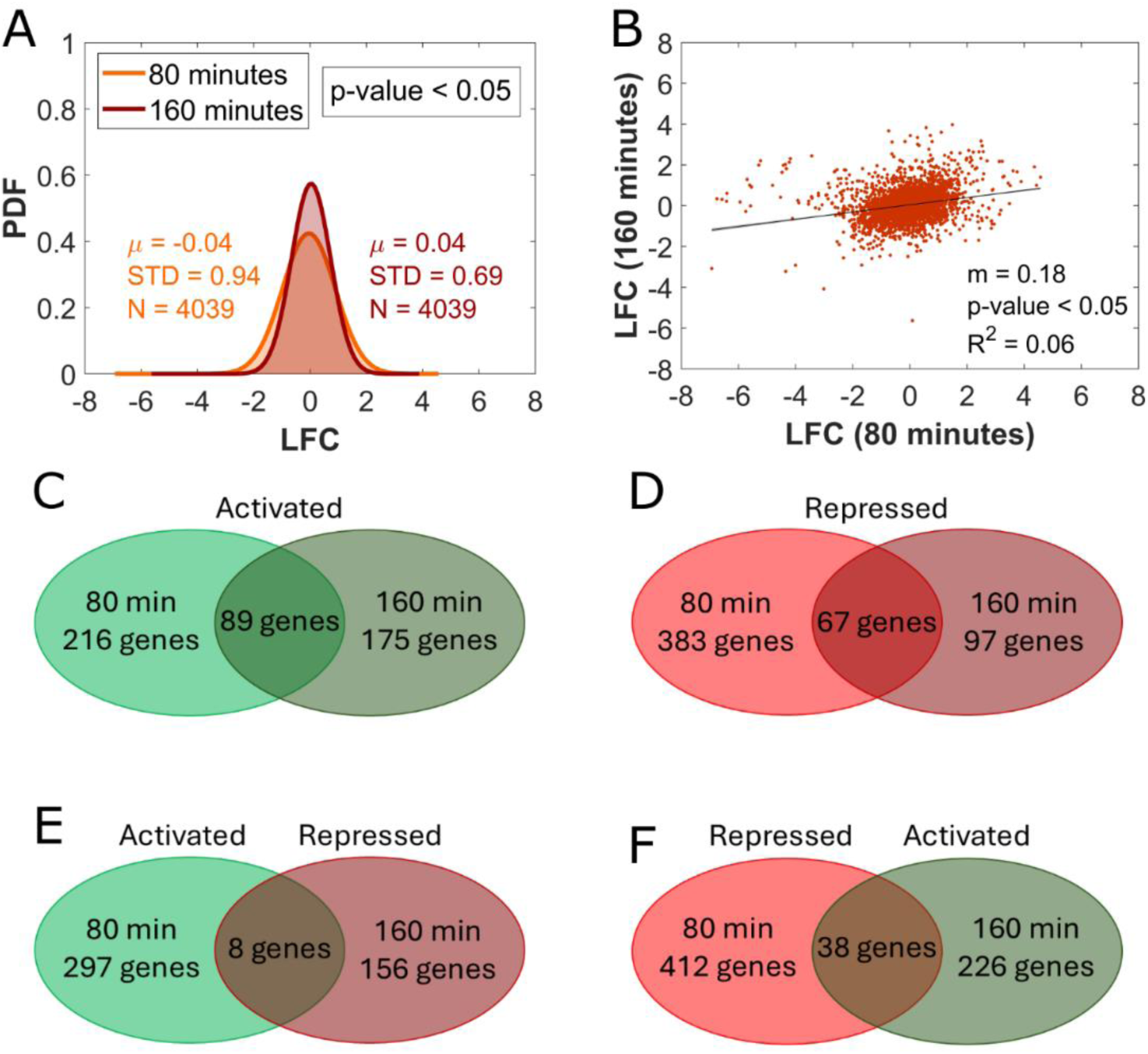
Time-lapse effects of rifampicin on the transcriptome. **(A)** Probability density functions of single-gene response strengths (Log_2_(fold change), i.e., LFC) relative to the control at 80 and 160 minutes after adding rifampicin, respectively. µ, STD, and N stand for mean, standard deviation and number of genes of the distribution, respectively. The p-value is from a two-sample t-test assessing if the two distributions are from independent random samples from normal distributions with equal means and equal, but unknown, variances. For p-value < 0.05, the t test statistics rejects the null hypothesis that the two distributions have equal means, at the 5% significance level. **(B)** Scatter plot of each gene’s response strength at 80 versus at 160 minutes. Shown is the best fitting line, its inclination (m), coefficient of determination R^2^, 95% confidence bounds (small shadow areas), and p-value under the null hypothesis that the line is horizontal. **(C)** Venn diagram of the number of genes activated under rifampicin at 80, 160, or both 80 and 160 minutes. **(D)** Same as (D) but for repressed genes. **(E)** and **(F)** are numbers of genes activated at 80 but repressed at 160 minutes, and vice-versa, or both, respectively.

Meanwhile, Figure 3B confirms that there is a significant correlation, at the single-gene level, between how each gene differs from the control at 80 and at 160 minutes, indicating that many changes occurring at 80 minutes remain at 160 minutes.

To further characterize the GRN behavior, we next focused on differentially expressed genes (DEG) (Methods Section 4.4). We classified DEG as: (i) activated if LFC > 1 (P-value < 0.05); (ii) repressed if LFC < -1 (P-value < 0.05); and (iii) non-responsive (i.e. non-DEG)), if -1≤ LFC ≤1 and/or P-value ≥ 0.05. Noteworthy, since some genes were removed from the analysis (Methods Section 4.4), the same gene cohort can differ in size between 80 and 160 minutes.

We found both positive and negative *early transient* (i.e., affected at 80 but not at 160 min)*, early persistent* (i.e., affected at 80 and at 160 min) *and late* (i.e., affected only at 160 min) DEGs. Also, some DEGs were early active/repressed, but switched to repressed/active at the late stage, respectively (Figures 3C, 3D, 3E, and 3F). Given this behavioral diversity, we hypothesized that several regulatory mechanisms were influential. In the next sections, we searched for such mechanisms.

### 3.4 Nucleotide changes in promoter regulatory regions can alter responses to rifampicin

DNA sequences with core promoters and operator sites (jointly referred to as ‘promoters’) largely control transcription initiation kinetics. Specifically, they influence RNAP-promoter association and dissociation kinetics (33, 34, 70, 71), open complex formation (72), and RNAP-promoter escape (73, 74) (Figure 4A). We investigated if promoter sequences affect the changes in RNA production rates caused by rifampicin.

**Figure 4.**
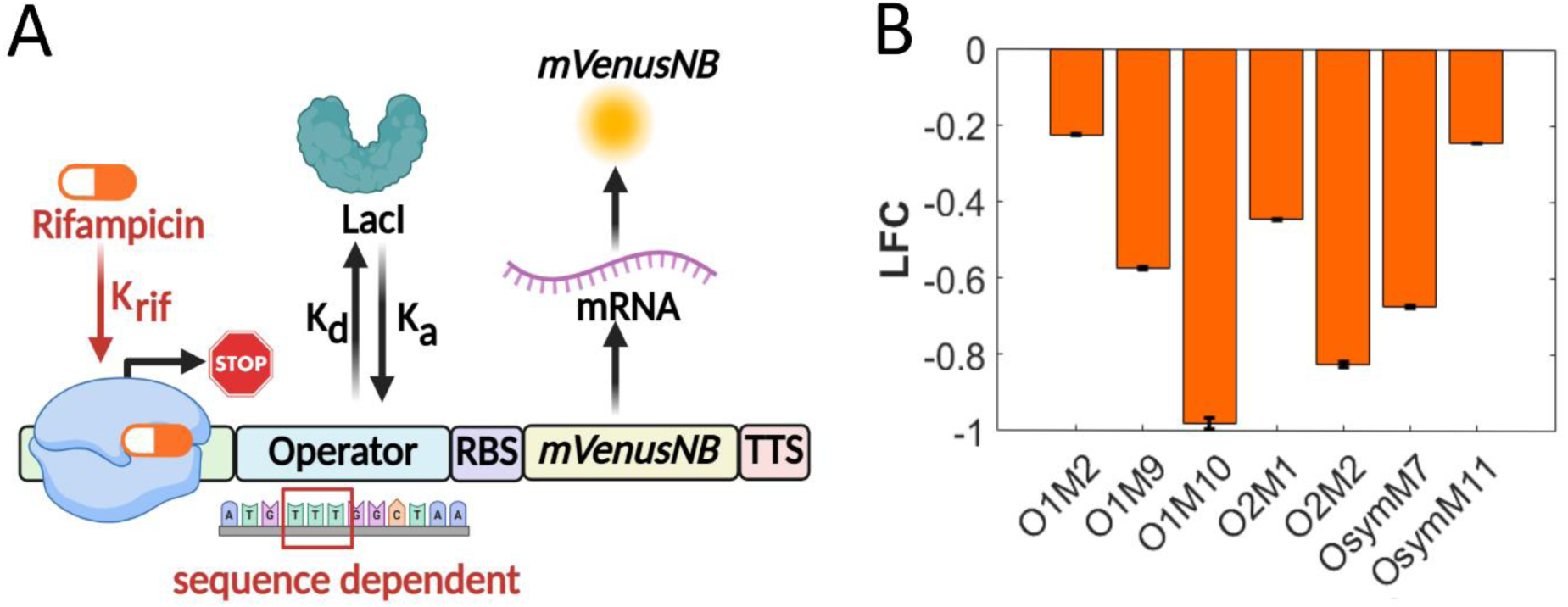
Effects of mutations in the operator region of a promoter on how rifampicin influences transcription, assessed using a strain library of chromosome-integrated Lac promoter mutants differing in LacI’s association and disassociation rates. **(A)** Illustration of a promoter with an RNAP binding region (core promoter) along with a binding site (operator) for the repressor LacI. The binding site sequence, which differs between strains, influences the binding (k_a_) and unbinding (k_d_) kinetics of LacI. Also shown is the mVenusNB coding region and corresponding RNA and fluorescent protein. TTS is the transcription termination site. **(B)** mVenusNB protein expression levels of each strain relative to the control condition measured by flow-cytometry (Methods section 4.5), 40 minutes after adding rifampicin. The error bars were calculated using the standard error of the mean (SEM) from 3 biological replicates and the propagation of uncertainty method.

To test this, it is necessary to have a set of genes that differ only in the promoter sequence. Since there is no natural set of genes with such property, we instead used a strain library (kind gift from the Elf lab) of cells with chromosome-integrated synthetic promoters pFAB120 with single LacI repressor binding sites that differ by a few nucleotides (Methods section 1). Shortly, we subjected cells to IPTG and, after transcription was activated and protein expression levels were stable (50 minutes after introducing IPTG), we next introduced rifampicin and compared the reductions in expression levels between strains (90 minutes after introducing IPTG). For this, we calculated the ratio between protein levels at 90 minutes (relative to the levels at 50 minutes) of each strain. We found that this ratio differs between strains, as expected if the promoter sequences influence how much rifampicin reduces transcription rates (Figure 4B).

### 3.5 Input TFs affect the early genome-wide response to rifampicin

Non-lethal genome-wide stresses can propagate in the GRN of E. coli via, e.g., transcription factors (TF) (24, 75–77). We investigated if TFs affect the GRN responsiveness to rifampicin, i.e. if TFs can alter the responsiveness of their output genes to be positive or negative depending on their regulatory effects (repressions or activations, available in RegulonDB (23)).

Using the methodology in (24), we considered the number of input TFs on each gene, k_TF_, and the regulatory effect of each TF (activation or repression, i.e. their individual TF bias). This information was obtained from (23). For each gene with more than one input TF, we estimated the ‘input TFs cohort bias’, *B*, by, first, assigning each repressor TF a ‘-1’, and each activator TF a ‘+1’ (if unknown, it was set to ‘0’). Then, we calculated the sum of these individual biases to estimate B, which is an approximate regulatory effect of the set of TF inputs (exemplified in Figure 5A). Finally, we separated the genes into cohorts according to their *B*.

**Figure 5.**
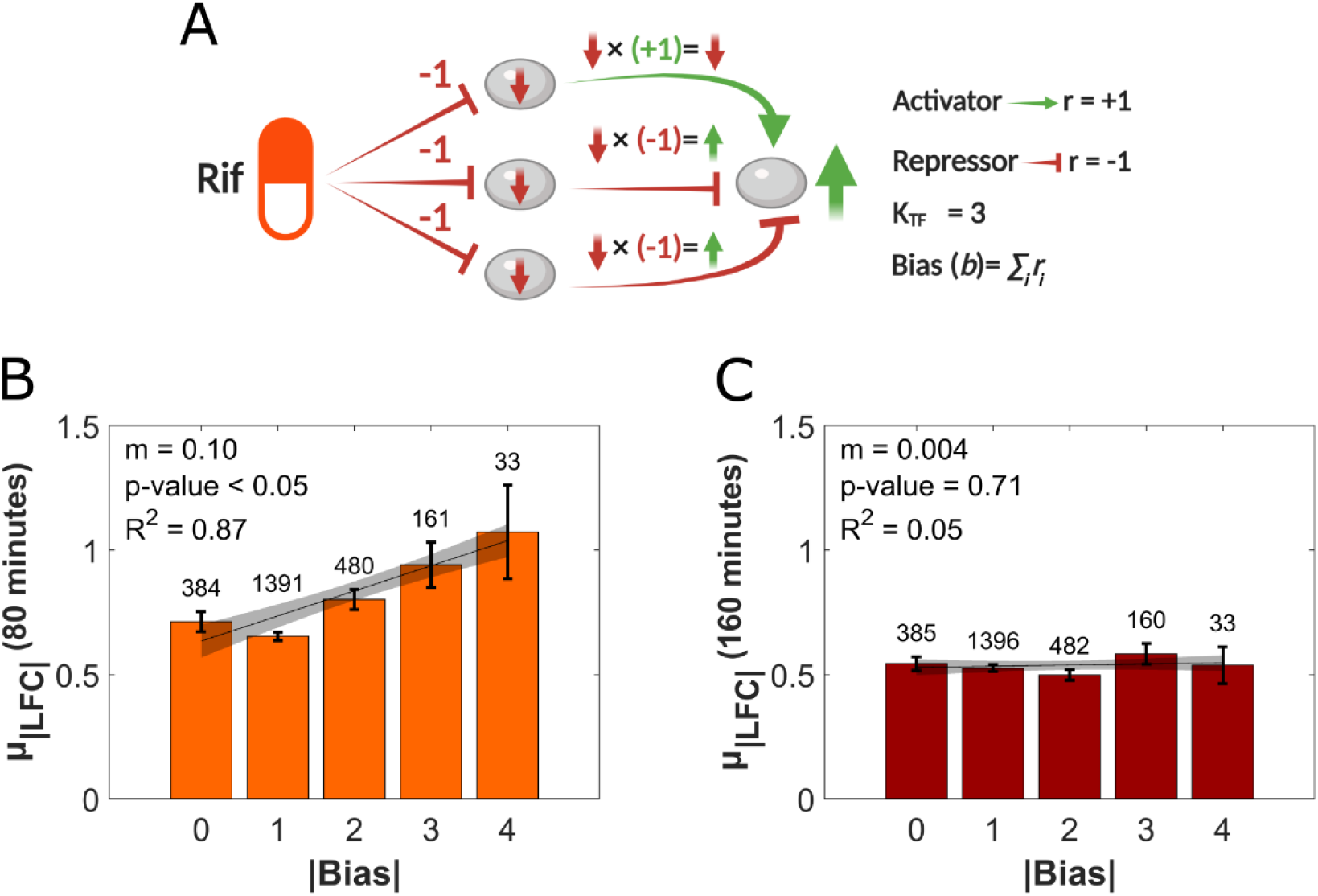
TFN and the early and late responses to rifampicin. **(A)** Illustration of the expected effects of rifampicin on an example gene subject to a set of 3 input TFs, whose regulatory role (r) consists of two TFs being -1 (repressions) and one being +1 (activation). If unknow, *r* would be set to 0 (not exemplified). The overall sum of the biases is, thus, +1. On average, we expect that rifampicin should first repress each of the input TFs, which in turn should cause the output to, eventually, be activated. **(B)** and **(C)** Average absolute LFCs, µ_|LFC|_, of gene cohorts as a function of their absolute bias |Bias| of their set of TF inputs at 80 and 160 minutes, respectively. The best fitting lines and 68% confidence boundaries (CB, red) were obtained using FITLM (MATLAB). p-values were obtained using the null hypothesis that the data is best fitted by a horizontal line. P-values are not rejected at 5% significance level. Also shown are the coefficients of determination (R^2^) and error bars (standard error of the mean, SEM). The numbers on top of the bars are the number of genes of the cohort.

Using this approximative model, we studied if the genes’ B correlates with the corresponding mean LFC, µ_LFC_. However, the results were inconclusive, statistically, since the p-value of the linear fit was higher than 0.05. This is likely because the error bars are large, potentially due to lack of sufficient data (Supplementary Figure S3A and S3B).

Therefore, we instead analyzed if the mean *absolute* LFCs, µ_|LFC|_, correlate with the mean *absolute* biases, |Bias|. From Figure 5B, µ_|LFC|_ is positively correlated with |Bias| at 80 minutes, with p-value < 0.05 and R^2^ > 0.85. Thus, the TFN logic and topology influence the early GRN response to rifampicin, likely being a major cause for transforming the initial genome-wide repressions into both repressions and activations, at later stages.

Finally, the correlation is not detectable at 160 minutes (Figure 5C). This is expected. If the genes change less after 80 minutes, they will cause less changes in their TF-neighbor genes as well.

Supplementary Figures S1A and S1B show that the conclusions hold when considering only ‘Activated’ or ‘Repressed’ genes, as classified in Results Section 2.3. Thus, from here onwards, we include all genes in our analysis, so that the conclusions are independent from the classification. Finally, since k_TF_ correlates positively with µ_|Bias|_ (Supplementary Figure S2A) (24), µ_|LFC|_ also increases with k_TF_ (Supplementary Figure S2B).

Finally, we plotted the LFCs of genes producing TFs against the LFCs of the genes that they regulate, as a function of the path length *L* between them, for *L* = 1, 2 (illustrated in Supplementary Figure 5A). We found positive correlations between the absolute LFCs of directly linked genes (L = 1). As expected, there is still correlation for L = 2, but weaker, both at 80 and 160 minutes. Overall, the results show that the transcriptome is TFN-dependent and, thus, GRN-dependent.

### 3.6 Neighboring genes display relatively similar responses to rifampicin

Evidence suggests that RNAPs can continue sliding in the DNA after transcribing a gene, potentially interfering with and correlating neighboring gene activities (78–80). Similar correlations can also be generated by, e.g., DNA supercoiling (65, 81).

We hypothesized that these or similar phenomena could cause neighboring genes to exhibit correlated response strengths to rifampicin. A similar observation has been reported for genes in the same operon (13). Here, we test if a similar phenomenon occurs between neighboring genes that are not in the same operon.

To test it, we considered the first and last gene of each operon (including operons with only 1 gene) (23) (Figure 6A). Next, we obtained the absolute differences between the response strengths (|ΔLFC|) of two genes not in the same operon, as a function of the distance ‘*D*’ between them (in number of operons, including operons with only 1 gene).

**Figure 6.**
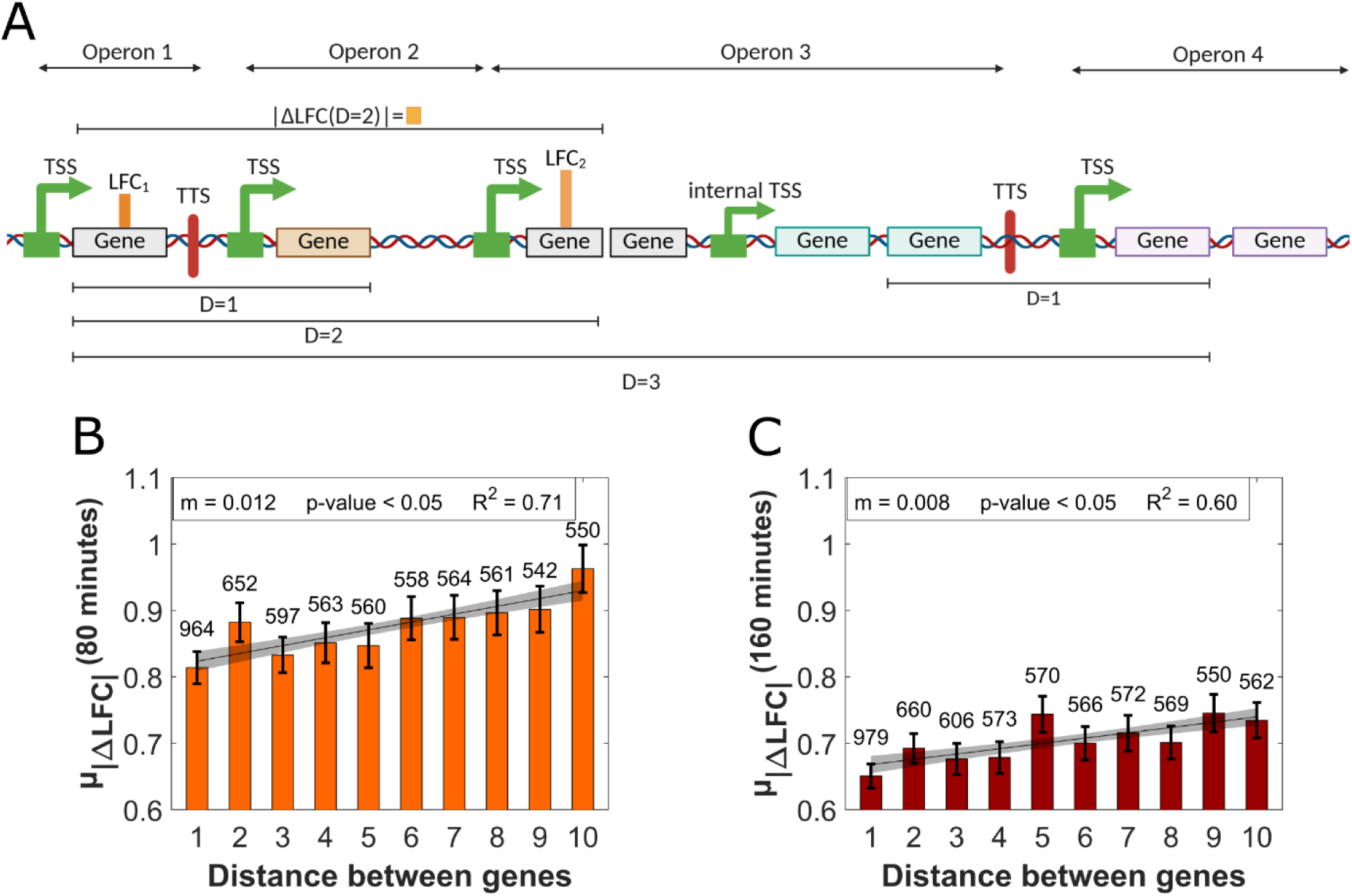
Differences in genes’ response strengths to rifampicin as a function of the distance between them (in number of genes). **(A)** Illustration of which pairs of genes were considered and how the distance and the absolute difference in response strengths (|ΔLFC|) as a function of the distance D were calculated. This illustration is adapted from one in (13). **(B)** and **(C)** are the average differences between the absolute response strengths of two genes (µ_|ΔLFC|_) as a function of the number of genes in between them at 80 and 160 minutes, respectively. Also shown are the best fitting lines and their inclination *m*, the coefficient of determination, R^2^, the 95% CBs (red shadows), and the p-value under the null hypothesis that the line is horizontal. The y axis does not start at 0 for easier visualization of the differences. Error bars are the standard error of the mean (SEM).

From Figures 6B and 6C, µ_|ΔLFC|_ increases with *D*. In detail, it increases significantly until D = 5 and then largely saturate after that. The increase from D=1 to 5 shows that the early and late response of genes to rifampicin is influenced by its neighboring genes.

Finally, Figure 6C (160 minutes after adding rifampicin) shows weaker differences and, thus, weaker m and R^2^ values. This is in line with the finding above that the RNA numbers differ less from the control at that time.

### 3.7 Gyrase levels decrease while genes sensitive to positive supercoiling buildup respond differently to rifampicin

Evidence suggests that rifampicin interferes with Gyrase (82). We thus measured Gyrase abundance (Methods Section 4.1), as it modulates positive DNA supercoiling buildup (Figure 7A). GyrB-YFP levels were higher than the control, but GyrA-YFP’s were lower (Figure 7B). Thus, Gyrase levels at 80 min are likely reduced by rifampicin.

**Figure 7.**
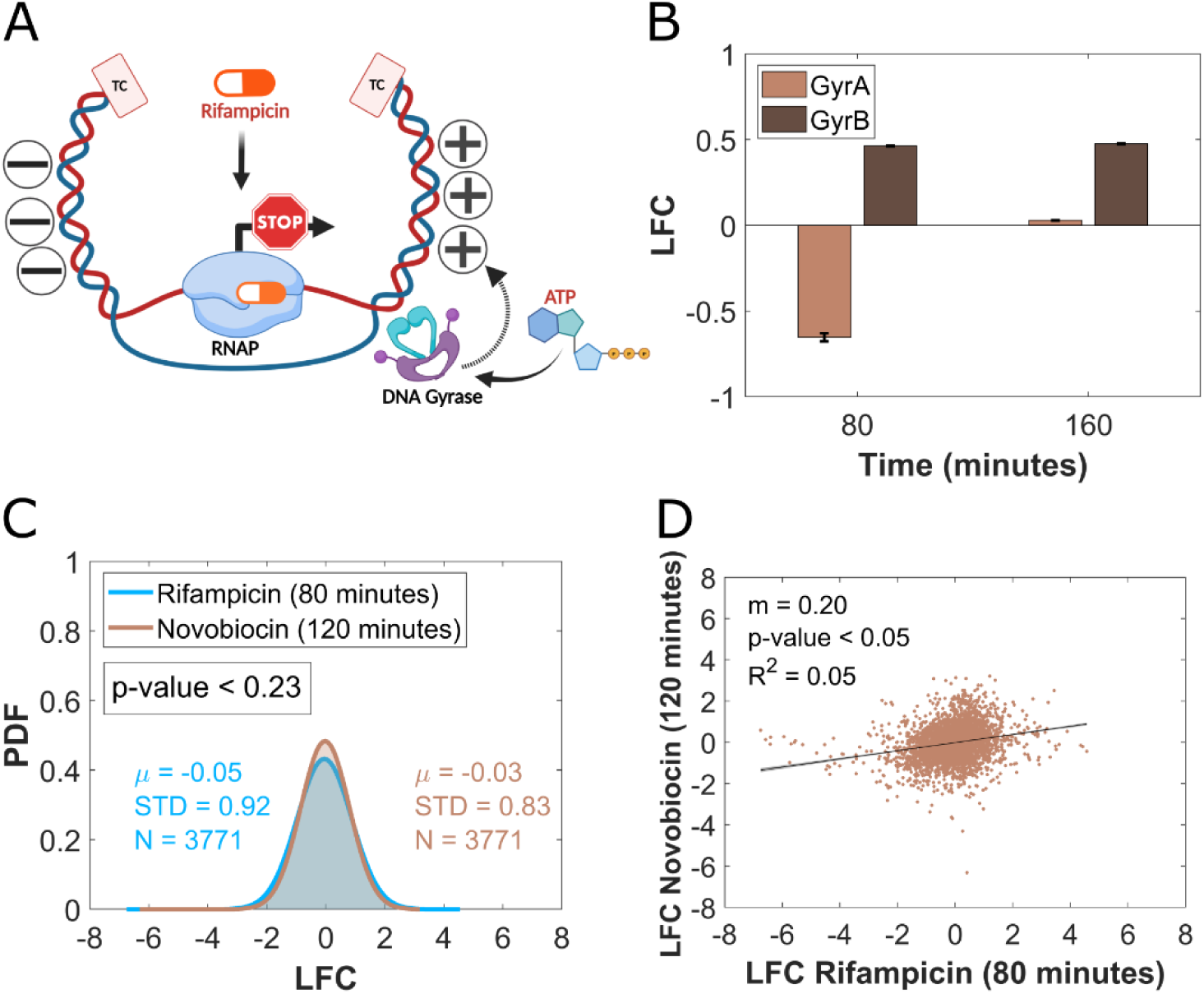
Sensitivity to positive DNA supercoiling buildup. **(A)** Generation (by RNAP) and maintenance (by DNA Gyrase) of positive DNA supercoiling in a topological domain, within two topological constraints (TC). Also shown is the interference of rifampicin on the generation of positive (+) and negative (-) DNA supercoiling, ahead and behind the elongating RNAP, respectively. **(B)** LFC (Log_2_ fold changes) of the mean single cell fluorescence levels of the two subunits of Gyrase (GyrA-YFP and GyrB-YFP), respectively, in cells subject to rifampicin relative to cells in standard growth conditions at 80 and 160 minutes after adding rifampicin. Error bars are calculated using the error propagation method. Error bars are the standard error of the mean (SEM) from N=3 biological replicates and the propagation of uncertainty method. **(C)** Probability density functions of the single-gene response strengths (LFC) relative to the control after adding novobiocin and after adding rifampicin, respectively. µ, STD, and N stand for mean, standard deviation, and number of genes, respectively. The measurements after adding Novobiocin were performed 40 minutes later than after adding rifampicin (Methods 4.3), in line with a slower permeation rate of Novobiocin, compared to rifampicin, of *E. coli* cell walls (14). The p-value is from a two-sample t-test assessing if the two distributions are from independent random samples from normal distributions with equal means and equal, but unknown, variances. For p-value < 0.05, the t test statistics rejects the null hypothesis that the two distributions have equal means, at the 5% significance level. **(D)** Scatter plot of the genes response strength (LFC) at 80 minutes to rifampicin and at 120 minutes to Novobiocin to account for different intake and response timings to the ABs (14). Shown also is the best fitting line, inclination (m), coefficient of determination R^2^, 95% confidence bounds (small shadow areas), and p-value under the null hypothesis that the line is horizontal.

Given this, we tested if genes sensitive to positive supercoiling buildup (PSB) respond to rifampicin distinctively from other genes. To test, we analyzed the LFCs of the 1215 genes identified previously as being more sensitive to PSB, due to their distinct responsiveness to Novobiocin (14), an AB that hampers Gyrase functioning (83). We found that their LFCs distribution at 80 min under rifampicin differs from other genes (t-test p-value < 0.05). This suggests that lowering Gyrase concentration (which likely increases PSB) significantly affects these genes’ responsiveness to rifampicin.

Contrarily, GyrA-YFP and GyrB-YFP levels suggest that Gyrase levels are similar to the control at 160 minutes, again limited by GyrA numbers (Figure 7B). In agreement, the LFC distribution of PSB sensitive genes subject to rifampicin does not differ from other genes (t-test p-value of 0.26).

### 3.8 Sensitivity to Positive Supercoiling Buildup correlates with the early response strength to rifampicin

Given that Gyrase levels were transiently reduced, we compared the LFCs of PSB sensitive genes, at 80 minutes under rifampicin with the LFCs of the same genes at 120 minutes under Novobiocin.

The difference in measurement times is due to a difference in how long these ABs take to cause an effect. While rifampicin alters RNA numbers in ∼10 minutes by blocking RNAP from escaping promoters (25, 26), Novobiocin interferes with Gyrase by blocking ATP binding, which is required to remove PSB (84). This causes PSB, which then affects RNA numbers. The changes become tangible ∼20-30 minutes after adding Novobiocin (14, 24).

We found that the two LFC distributions do not differ significantly (Figure 7C). Moreover, the LFCs of each gene under each of the ABs correlate positively (Figure 7D). This correlation is not due to a relationship between cell growth and gene expression since we calculated LFCs using a control at the same point in time.

Instead, the correlation is in line with the transient decrease in Gyrase levels (Figure 7B). Moreover, by affecting PSB, Novobiocin could also affect (among other steps) the same step as rifampicin, promoter escape. This could further explain why neighboring genes exhibited relatively closer response strengths to rifampicin than distanced ones.

### 3.9 σ^32^, σ^38^, and σ^70^ influence the genes responsiveness to rifampicin

RNAPs require σ factors (85) to recognize promoter sequences (23) (Figure 8A). Since cells have less RNAPs than σ factors, changing their numbers alters the transcription rate of promoters with different σ factor preferences (34, 86–88). E.g., increasing the numbers of one σ factor will increase the transcription rates of its output genes and decrease the transcription rate of all other active genes (89).

**Figure 8.**
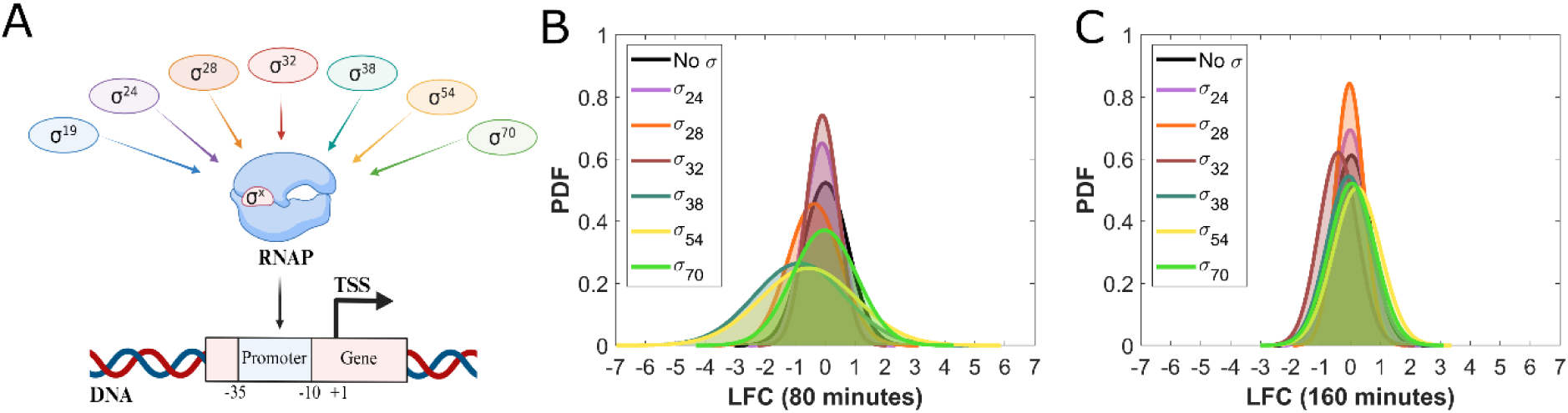
σ factors and the response to rifampicin. **(A)** Illustration of the mechanism of σ factor regulation, where some promoters can only be transcribed after the binding of a specific σ to RNAP. **(B)** and **(C).** Probability density functions of the single-gene response strengths relative to the control (LFC) of genes whose promoters are recognized by a specific σ factor after 80 minutes and after 160 minutes of rifampicin exposure, respectively. All genes are considered, along with their nearest upstream promoter, irrespective of their positions in the operons. Mean, standard deviation, and number of genes of each distribution shown in Supplementary Table 1, along with the p-values *of two-sample t-tests to assess whether* two distributions are from independent random samples from normal distributions with equal means and equal, but unknown, variances. ‘No σ’ stands for ‘no known σ’.

Since σ^32^ might affect the responsiveness to rifampicin (90), we compared genes’ responses to rifampicin as a function of the σ factors. Genes with dual σ factor regulation (less than 400 (23)) were included in the cohorts of both σ factors.

At 80 minutes, both σ^38^ and σ^70^ and their output genes behaved (statistically) distinctively from other genes (Figure 8B and Supplementary Table S1). However, while both mean and STD of the distribution of LFCs of the σ^38^ regulated genes differ from other genes, for genes regulated by σ^70^, only the variability differs.

At 160 minutes, only σ^32^ and its output genes differed from the control (Figure 8C and Supplementary Table S2), in agreement with (90). Overall, these results support the hypothesis that the early response to rifampicin is influenced by the GRN.

### 3.10 (p)ppGpp affects gene responses to rifampicin

If rifampicin affects (p)ppGpp, it could indirectly affect RNAP-promoters selectivity (91) (Figure 9A). We tracked *spoT*, one of two genes (the other being *relA*) responsible for (p)ppGpp production (20). Also, we assessed if (p)ppGpp-regulated genes responded differently to rifampicin (308 genes according to RegulonDB Vs. 13.0 (23)).

**Figure 9:**
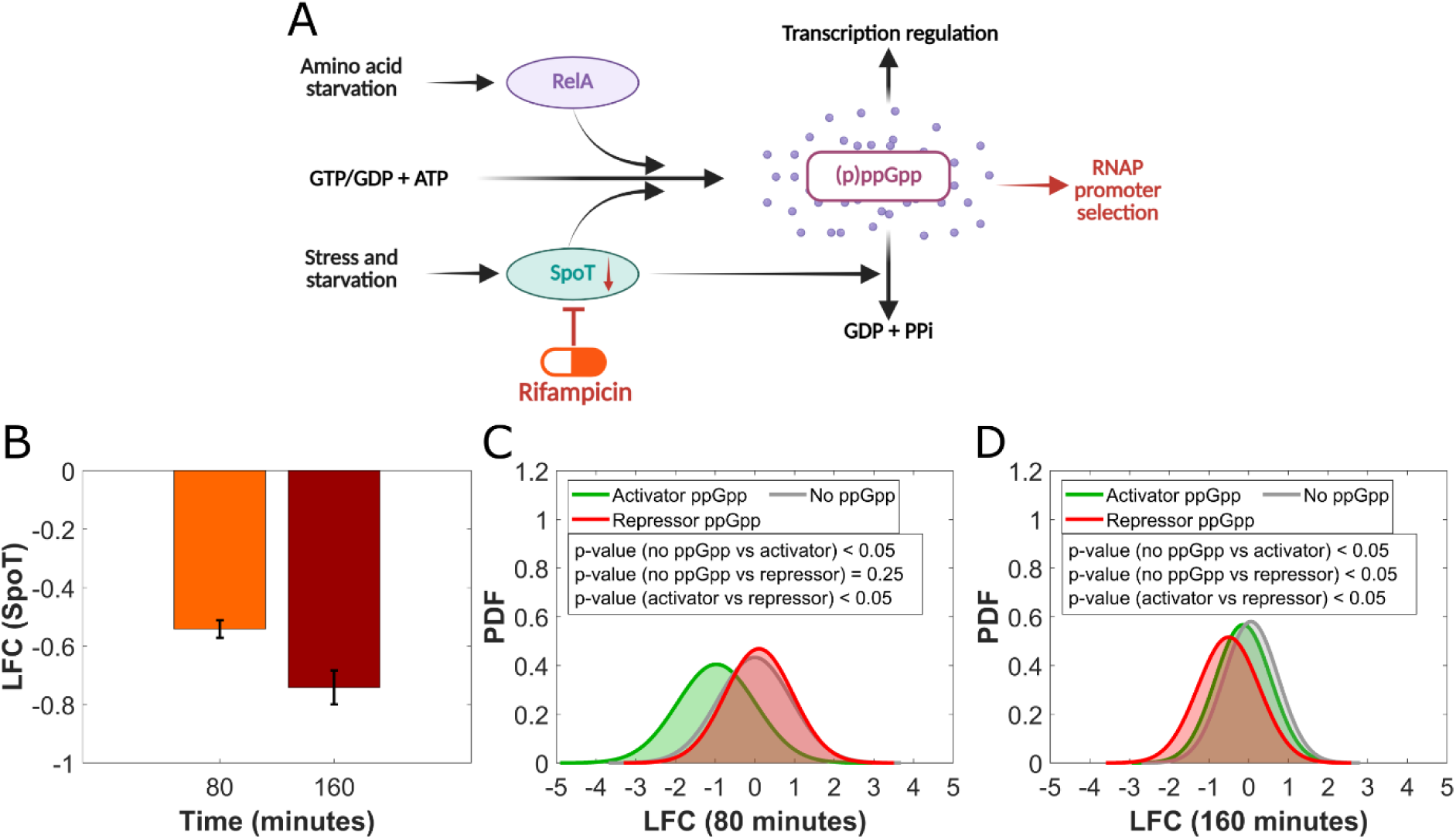
(p)ppGpp influence of the global response to rifampicin. **(A)** Illustration of the mechanism of (p)ppGpp regulation, with the two genes responsible (relA and spoT) influencing how RNAP selects promoters. Adapted from (92). **(B)** SpoT expression levels relative to the control (measured by SpoT-YFP) at 80 and 160 minutes. Error bars calculated using the standard error of the mean (SEM) from 3 biological replicates and the propagation of uncertainty method. **(C)** and **(D)** Distributions of LFCs of genes known to be activated (µ = -0.97, STD = 0.96, N = 182), repressed (µ = 0.10, STD = 0.85, N = 99), and not regulated by (p)ppGpp (µ = 0.00, STD = 0.92, N = 3764), at 80 minutes and 160 minutes, respectively. Also shown are the p-values of two-sample t-tests to assess if the two distributions are from independent random samples from normal distributions with equal means and equal, but unknown, variances. For p-value < 0.05, the t test statistics rejects the null hypothesis that the two distributions have equal means at a 5% significance level.

We found that SpoT levels were lower than the control at 80 and at 160 minutes (Figure 9B). In agreement, most genes upregulated by (p)ppGpp were more repressed, while most genes downregulated by (p)ppGpp were more activated than control genes (Figures 9C and 9D).

### 3.11 The global regulators Fnr, Fis, LexA, and MarA and their regulons behaved differently from the control

A group of 20 TFs controls ∼30% of *E. coli* genes (23). Many of these ‘global regulators’ (GR) are involved in stress responses. We investigated if these GRs respond to rifampicin and if the genes that they control also react accordingly. For this, we used a strain library of single-copy plasmids with 16 out of the 20 promoters of the GRs of interest controlling GFP expression [89] (the remaining 4 are not available in the library). Since the GRs activate some genes but repress others, we studied *absolute* log_2_ RNA fold changes under rifampicin.

Supplementary Tables S3 and S4 show the responses of each GR and the corresponding cohort of output genes at 80 minutes as well as at 160 minutes. From there, we searched for cases where the GR and output genes exhibit consistent behaviors. We found four cases where both the GR as well as its output genes (regulon) differed from the control. Specifically, fnr and fis at 80 minutes, and lexA and marA at 160 minutes.

Relevantly, the GRs biofunctions are consistent with their presence assisting *E. coli* in adapting to rifampicin in the absence of resistance mechanisms. Specifically, they are involved in chromosomal organization (Fis), DNA damage response (LexA), anaerobic respiration (Fnr) and multiple AB resistance (MarA), respectively (23). The overexpression of MarA is particularly significant since this gene activates multidrug efflux processes (93–96) and reduces membrane permeability (97), which could explain the smaller difference in RNA levels relative to the control at 160 minutes.

To support these results, we also analyzed the single-cell expression levels of Fnr, Fis, LexA, and MarA using flow-cytometry (Supplementary Figure S3). We also considered SoxS, due to its role in AB resistance [90]. In agreement with the RNA-seq data, all mean expression levels under rifampicin were lower than the control at 80 and 160 minutes (except for Fnr at 80 min and LexA at 160 min).

### 3.12 DNA strands, promoter AT-Richness and P-distances, sRNAs, RNAP-Promoter binding rates did not affect the response to rifampicin

We assessed if other features, in addition to the ones above, could be associated with the genome-wide selective responses to rifampicin.

First, of the 4748 *E. coli* genes (RegulonDB 13.0, (23)), 2349 genes (49.5%) are in the forward strand while 2399 (50.5%) are on the reverse strand. Although the difference in numbers is small, these two cohorts could be influenced differently by replication, which occurs in only one direction. If so, this could then affect the average responsiveness to rifampicin. Thus, we assessed if the genes in one strand were more affected by rifampicin than the other genes. For this, we compared the LFC distributions of genes in different strands. However, we did not find differences (statistically), either at 80 or at 160 minutes.

Second, we searched for correlations between the genes’ LFC and their promoter sequence (AT-richness and p-distance, respectively (Methods section 4.8)). We did not find correlations either at 80 or at 160 minutes.

Third, we investigated regulatory non-coding RNAs (sRNAs). We considered genes positively and negatively regulated by sRNAs, respectively (23), but did not find differences in their LFC distributions compared to the genome-wide distribution, either at 80 or at 160 minutes.

Finally, we considered that transcription initiation is a multi-step process that includes closed and open complex formations, as well as promoter escape (99) and is followed by elongation (for a review of this process, see (100)). Evidence suggests that, while rifampicin can bind RNAP at any point prior to elongation, it only affects the rate of escaping promoters rather than, e.g., its promoter binding affinity (25). If this is true, its effects on individual genes should not correlate with the effects of reduction RNAP intracellular concentrations. These, by reducing free floating RNAP concentrations (101), should affect only closed complex formations, whose durations are not expected to correlate with RNAP promoter escape rates. In agreement, from data of LFCs when reducing RNAP intracellular concentrations (24), we did not find them to correlate with the LFCs caused by rifampicin (Supplementary Figure 4).

### 3.13 Orthologous genes of *E. coli* and *M. tuberculosis* have correlated response strengths to rifampicin

*M. tuberculosis* is evolutionarily distant from *E. coli* (different Phyla that diverged long ago) (102). Also, it has higher GC content (∼65%) (103) than *E. coli* (∼50%) (104) and its cell walls differ structurally (105). This latter difference is particularly relevant, since two reasons for *E. coli* to have high survivability against rifampicin may be lesser membrane permeability along with more efficient membrane efflux pumps than *M. tuberculosis* (106). Finally, the physiologies of these species also differ. While *M. tuberculosis* is a pathogen, grows slowly, and persists in host tissues (107), *E. coli* grows fast and lives in guts without harming them (108).

We limited the analysis to *M. tuberculosis* since we did not find RNA-seq data on non-lethal rifampicin concentrations at similar time points for any other species. Data on *M. tuberculosis* (six strains) subject to non-lethal rifampicin concentrations was obtained in the NCBI GEO database (GSE292409). It informs on TPMs (Transcripts Per Million). Using Mycobrowser (https://mycobrowser.epfl.ch/ as of June 28th, 2024) (109), we associated each locus and gene identifier with their corresponding TPM, with the aim of comparing with our *E. coli*’s TPMs (Methods section 4.4).

Before that, we first compared each gene’s log₂(TPM) between two of the six strains of *M. tuberculosis*. The strong correlation observed (Supplementary Figure S5) suggests that the measurements are robust. As such, this correlation can be used as an estimation for a maximum correlation that could be expected when comparing log₂(TPM) of different species. Comparing other pairs of the six *M. tuberculosis* strains gave similar results.

Next, we searched for orthologous gene pairs of *E. coli* and *M. tuberculosis* (i.e. genes evolved from a common ancestral gene). For this, we retrieved the IDs of each *E. coli* gene and queried the NCBI Gene database using the rentrez R package (110) in order to identify orthologs genes in *M. tuberculosis*. We found 567 orthologous pairs.

Next, we obtained the corresponding log₂(TPM)s under non-lethal rifampicin concentrations of those genes for both species (with the available data being at 80 and 60 minutes after adding rifampicin for *E. coli* and *M. tuberculosis*, respectively). We found that the TPMs of orthologous genes are correlated (Supplementary Figure S6 and Supplementary Table S5), albeit weaker than between the two *M. tuberculosis* strains (as expected).

Given that orthologous genes of these evolutionary distant species respond similarly to rifampicin, we argued that both species are exerting control over which genes respond and how strong are their responses to non-lethal rifampicin stress. Further, the similarity in the genes’ responsiveness suggests that the two species are triggering similar (likely beneficial) phenotypic changes, associated with the biofunctions regulated by these genes.

3.14 Adjacent genes on the DNA exhibit similar response strengths in both *M. tuberculosis* and *E. coli*

We aimed to compare gene responses of *M. tuberculosis* and *E. coli* to rifampicin as a function of their regulatory mechanism (e.g. TFs and promoter sequences). Unfortunately, information on such mechanisms is unavailable for most species, including *M. tuberculosis* (e.g. on TFs). However, the chromosomal position of genes relative to the origin of replication is available for both species (NCBI). As such, the effects of distances in the differences between the genes’ response strengths to rifampicin can be compared.

First, we estimated distances between genes. Next, we obtained the average absolute differences in responses strengths to rifampicin (*µ_|log₂(TPM)|_*) for all pairs of genes, as a function of their distance in the DNA *D*, i.e. number of genes in between, for *D* ≤ 50. From Supplementary Figures S7 and S8, in both *E. coli* and all strains of *M. tuberculosis*, *µ_|log₂(TPM)|_* increases with *D* in a manner that is well fitted by an exponential function of the form: y = c + a×e^-bx^ (best fitting parameters in Supplementary Table S6). Specifically, all R^2^ and adjusted R^2^ values are above 0.85.

Moreover, we did not find significant differences in the parameter values of the best fitting curves of the different *M. tuberculosis* strains. This suggests that the method to compare curves is relatively precise. Noteworthy, while the values of *b* (equation growth rate) are similar between species, *a* and *c* (starting and saturation points, specifically) differ widely. We do not know why, but it may be due to differing total number of intracellular RNAs of the two species and/or differences in the RNA-seq methods of the studies.

Overall, since both species show similar TPM differences with the distance between genes, the underlying causes might be the same (potentially local positive supercoiling buildup, RNAP traffic and interference, or the replication machinery).

### 3.15 The features affecting gene responses are enriched in *E. coli* orthologs of *M. tuberculosis* genes

We explored whether *E. coli* orthologs of *M. tuberculosis* possess (more often than expected) features identified above as influential in the early response to rifampicin (e.g., sensitivity to PSB). In general, we identified how many *E. coli* genes are both orthologs (‘A’) and possess another feature (‘B’). Next, we estimated how many genes, out of a total of N genes, would have both properties, A and B (N_A∩B_) by random chance: *N_A∩B_=P(A∩B)×N = P(A)×P(B)×N*. Here, *P* stands for the fraction of genes with a given feature. We only considered *E. coli* genes for which there is LFC data at 80 minutes (4047 genes), which are 565 out of the 567 genes orthologs of *M. tuberculosis*.

Next, we estimated which difference (in number of genes) between empirical data and random chance should be considered significant. For that, we a property that was not influential in the response strength to rifampicin, which is the DNA strand where the genes are located (Results section 2.12). Thus, we calculated the difference in number of genes between strands for orthologous genes. From Supplementary Figure S9, the number of genes in the forward DNA strand according to the empirical data and the numbers expected by chance differ by ∼8% (the same difference is true for the reverse strand). From there onwards, we use this percentage as a “minimum difference’ in number of genes with a given feature, above which we consider the difference significant.

We next studied if the features for which we found evidence to be influencing early gene responses to rifampicin are common in *E. coli* orthologs of *M. tuberculosis* genes. From Figure 10, we found that, in all cases, the number of orthologous genes with each of those properties are higher than expected by chance, respectively.

**Figure 10.**
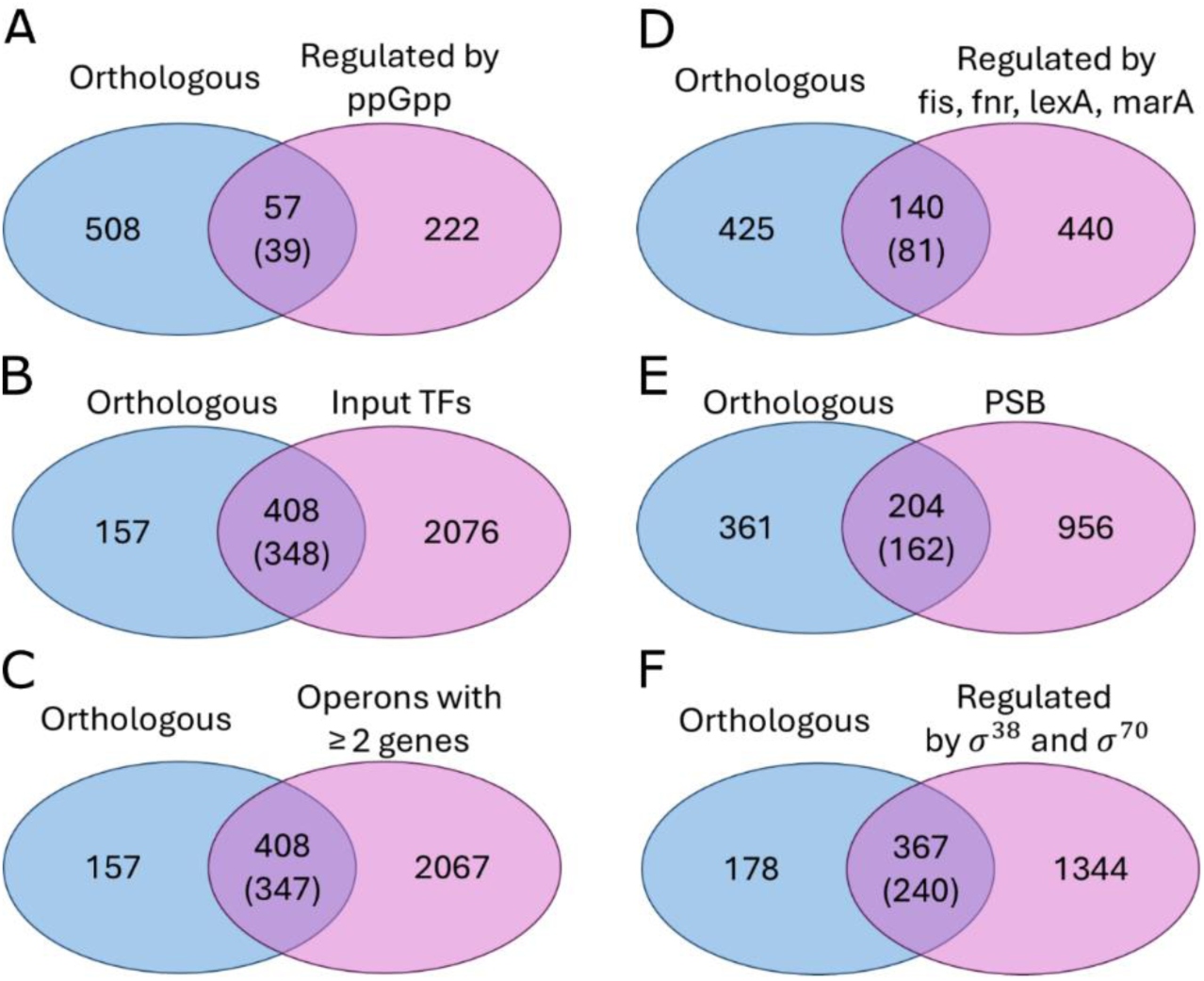
Venn diagrams of *E. coli* orthologs and *E. coli* genes with regulatory mechanisms affecting rifampicin response at 80 minutes. The intersecting areas show the number of genes with both properties. Inside the parentheses are the number of genes expected by random chance to have both properties. **(A)** genes regulated by (p)ppGpp; **(B)** genes with input TFs; **(C)** genes in operons with more than one gene; **(D)** genes regulated by Fnr, Fis, LexA, or MarA; **(E)** genes sensitive to PSB; and **(F)** genes regulated by σ^70^ and/or σ^38^.

Specifically, *E. coli* orthologs to *M. tuberculosis* have a larger than expected number of (p)ppGpp regulated genes (32% more than expected), genes with input TFs (29% more than expected), presence in operons with more than 1 gene (15% more than expected), regulation by Fnr, Fis, LexA *or* MarA (57% more than expected), sensitivity to PSB (20% more), and σ^38^ regulation (51% more).

Overall, the results suggest that not only orthologs exhibit similar responses to *M. tuberculosis* genes but that this is largely due to the features found above to influence those responses.

### 3.16 *E. coli* genes orthologous to *M. tuberculosis* do not have enriched promoter AT-Richness or P-distances. sRNAs regulation is slightly more enriched

Finally, we studied whether the features found to not influence early gene responses could be related, or not, with orthologs. Again, we considered 8% as the percentage corresponding to a “minimum difference’ below which the results are considered not differing.

From Supplementary Figure S10, the percentage of genes that are both orthologous and have higher than average AT richness is smaller than expected by chance. Further, the percentage of genes that are both orthologous and have higher than expected p-distance is lower than 8% (thus, not significant). Potentially, this could be explained by the significant role of σ^38^ regulated promoters (p distances correspond to preference for σ^70^). Finally, the percentage of orthologous genes that are regulated by sRNAs (263, i.e., 46%) also differs little from expected by chance (∼5% < 8%).

Overall, these results support our conclusions above that these features do not influence the response to rifampicin, either globally or of ortholog genes of *E. coli* and *M. tuberculosis*.

## 4. DISCUSSION

We studied the response of E. coli to non-lethal rifampicin stress, as the initial effects propagated through the GRN. We observed a strong, transient change in the transcriptome and found several components and features of the GRN causing it.

First, RNAP numbers increased. This could facilitate cells overcoming rifampicin effects, by increasing free-floating RNAPs and RNAP collisions to free rifampicin-trapped RNAPs at promoter regions. Contrarily, it may be energetically costly, given the need to synthesize more RNAPs and σ factors and having to cope with additional PSB with reduced Gyrase levels (14).

Second, changing a few nucleotides in promoter sequences affected rifampicin efficiency in reducing RNA production. E. coli could exploit this to tune each promoter. Contrarily, this might cause harmful asymmetries in the production of protein components or in pathways involving more than one protein. It can also disrupt single-cell protein numbers bimodalities (111).

Third, the TFN propagated the initial rifampicin perturbations, and the strength of the signals propagated differed with the number and regulatory logic (overall activation vs. repression) of input TFs of each gene. Potentially, there could exist a layer of genes detecting rifampicin (and/or signal filtering), followed by a layer of genes enhancing cell adaptation, particularly given that the RNA levels have changed widely between early and late responses. Contrarily, some of the propagated signals may have negative consequences, such as the deregulation of genes that otherwise were robust to rifampicin, potentially harming the cell and causing chaotic GRN dynamics.

Fourth, neighbor genes in the DNA, even in different operons, had comparatively similar responses to rifampicin. This could be due to PSB, which differs between topological domains (65), affecting the efficiency of rifampicin in hampering transcription initiation. In support, PSB sensitive genes differed in rifampicin responsiveness. The phenomenon may allow compartmentalize topologically the effects of rifampicin. If so, genes with beneficial, complementary functions or involved in the same processes could be organized in the same, less affected domain(s). Placing gene sets in more affected domains could also be beneficial, e.g. for synchronized shutdown. Contrarily, the phenomenon might harm multi-step processes.

Fifth, a few global regulators (σ factors and GRs) responded to rifampicin and changed their output genes accordingly. Future studies on the phenotypic changes caused could provide insight into how bacteria increase AB tolerance in the absence of resistance mechanisms.

Sixth, genes upregulated/downregulated by (p)ppGpp were significantly repressed/activated, respectively. In agreement, SpoT was repressed. This may assist cells avoiding transcription shutdowns and thus enhance cell repair, energy maintenance, and efflux pumping. Contrarily, in the long-term, it can hamper the regulation of cell processes requiring (p)ppGpp.

Seventh, genome-wide single-gene responses to rifampicin correlated with those to Novobiocin, albeit their MoAs differ (65). Potentially, a similar set of mechanisms drove the transcriptome in both cases. If so, findings reported here could apply to aminocoumarins and, potentially, other AB families, such as fluoroquinolones, as they also target Gyrase.

Finally, we studied how the findings could apply to an evolutionary distant species, M. tuberculosis. We found several similarities in how the genome and transcriptome of the two species when under rifampicin. First, the response strengths of orthologous genes of E. coli and M. tuberculosis correlate. This could be evidence for conserved rifampicin-responsive genes from a common ancestor, or that both species independently evolved similar phenotypic adaptations that are triggered by orthologs. In support of the first hypothesis, we found that all the features that we found to affect the gene responses in E. coli were enriched in orthologs of M. tuberculosis genes. In further support, the features that we did not finding evidence to affect gene responses, were not enriched in orthologs. Second, we found a similar regulatory phenomenon in M. tuberculosis. Namely, adjacent genes in the DNA also exhibited similar response strengths to rifampicin. This is evidence of a similar regulatory mechanism (1-dimensional signal propagation between neighbor genes), potentially used to improve survivability. Future studies of the GRN of M. tuberculosis, as in (112), could provide additional information.

In summary, our findings suggest that the response of E. coli’s GRN to rifampicin was affected by several local and global features, encoded in gene sequences, and also embedded in various layers of the regulatory structure. Some of the features (e.g., TF and other GRs numbers) are known to be able to adapt the cells to various conditions. This suggests that the response to rifampicin of the GRN of E. coli might be adaptive, even in the absence of resistance mechanisms. Moreover, other features (e.g., promoter sequences) are evolvable, supporting that E. coli’s response to rifampicin is evolvable. Future studies should be able to verify the positive and negative phenotypic roles of these features and find valuable applications.

Several questions remain unanswered particularly on how some of the features above are triggered and how does it benefits cell survivability. E.g., how do input TFs alter the effect of rifampicin on output genes? Does it depend on their mode of action? It would also be interesting to expand the study to assess the outcome of combinations of relevant features, i.e. studying the responses of genes combining two or more features (e.g. (p)ppGpp and σ38 regulation). Arguably, genes conducting the most valuable adaptations of the cell might

possess more than one response mechanism, to enhance responsiveness. Other genes could make use of requiring the activation of more than one mechanism in order to be responsive.

Finally, by dissecting natural genes’ rifampicin-response mechanisms, our findings could support engineering valuable synthetic circuits with scientifical and/or biotechnological applications. For example, the ability of promoter sequences to affect how much rifampicin obstructs their RNA production rate could be of use to tune the responsiveness of future AB responsive (dose-dependent) synthetic genetic circuits (e.g., genetic toggle switches, clocks, and signal filters). Also, the ability of TFs to propagate the effects of rifampicin could be used in the engineering of large synthetic genetic circuits. Our findings could also contribute to understanding how bacteria trigger transcriptional programs of AB tolerance (113), particularly by identifying genes involved, their trigger mechanisms, and whether they are unique to E. coli or common in other bacteria. This could have implications in the development of new ABs that, rather than targeting resistance, instead target the strategies of AB-survivability applied by AB susceptible species, which influences their ability to develop resistance.

## AUTHOR CONTRIBUTIONS

A.S.R conceived and directed the study. M.M.A. assisted in the planning along with R.J. M.M.A. executed most data analysis, assisted by R.J., A.M.A., and A.S.R. Meanwhile, A.A. executed the measurements. M.M.A., A.A., R.J., and A.S.R. interpreted and integrated the data. A.S.R, assisted by A.A. and M.M.A., drafted the manuscript. M.M.A. produced most figures assisted by A.M.A., R.J., and A.S.R. R.J. and A.M.A. produced a few figures. All text and figures were revised by all authors.

## FUNDING

Work supported by the Sigrid Jusélius Foundation [grant numbers 230181, 240191, and 250213] to A.S.R., Suomalainen Tiedeakatemia [to R.J.]; EDUFI [grant number TM-24-12142] to A.A and [grant number TM-25-12197] to M.A., and the Doctoral Programme of the Faculty of Medicine and Health Technology of Tampere University [to A.A]. The funders had no role in study design, data collection and analysis, publication, or preparing the manuscript. Funding for open access charge: Tampere University.

## ACKNOWLEDGEMENTS

We thank Johan Elf, Hiromi Imamura, and Robert Landick, for their generosity in sharing strains with us. We thank Vinodh Kandavalli, Cristina Palma, Ines Baptista, Vatsala Chauhan, and Howy Jacobs for valuable suggestions and information.

## CONFLICT OF INTEREST

The authors have no competing interests.

## Notes

### Competing Interest Statement

The authors have declared no competing interest.

