## Supplemental Figures and Tables for "Mechanisms shaping the transcriptome of E. coli to non-lethal rifampicin stress"

**Supplementary Figures**

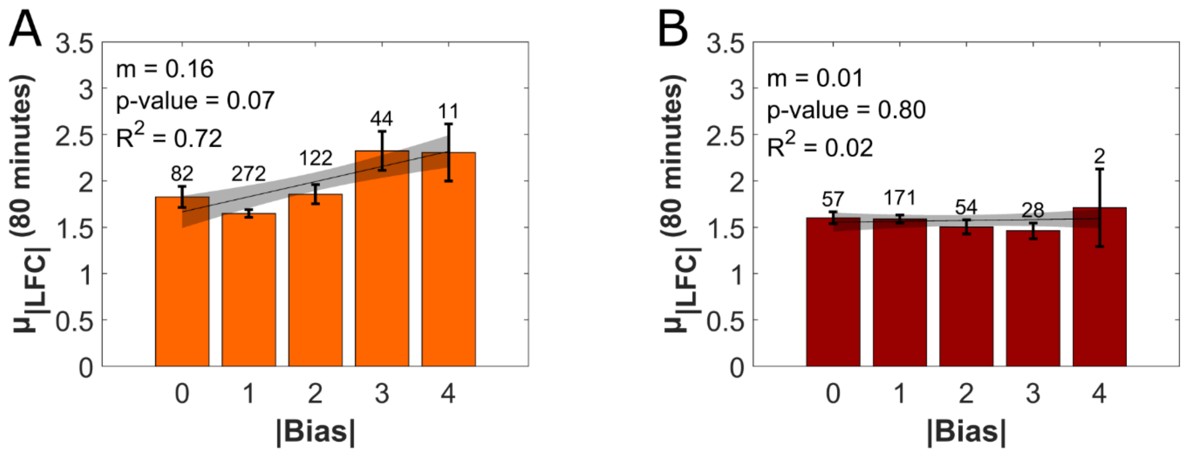


**Figure S1**: Genome-wide effects of the TFN on the **(A)** early and **(B)** late responses of the GRN of *E. coli* to rifampicin, when considering only DEG genes, i.e., genes whose LFC has a p-value < 0.05 & |LFC|>1. Shown are the average absolute LFCs, $\mu_{|LFC|}$ of genes whose set of TF inputs has a given mean absolute bias. The best fitting line and 68% confidence boundaries (grey) were obtained using the Fitlm function of MATLAB. Also shown are the coefficient of determination $R^2$, inclination of the line *m*, and p-value under the null hypothesis that the line is horizontal. The hypothesis is not rejected for p value < 0.05. On top of each bar is the number of genes of the cohort.

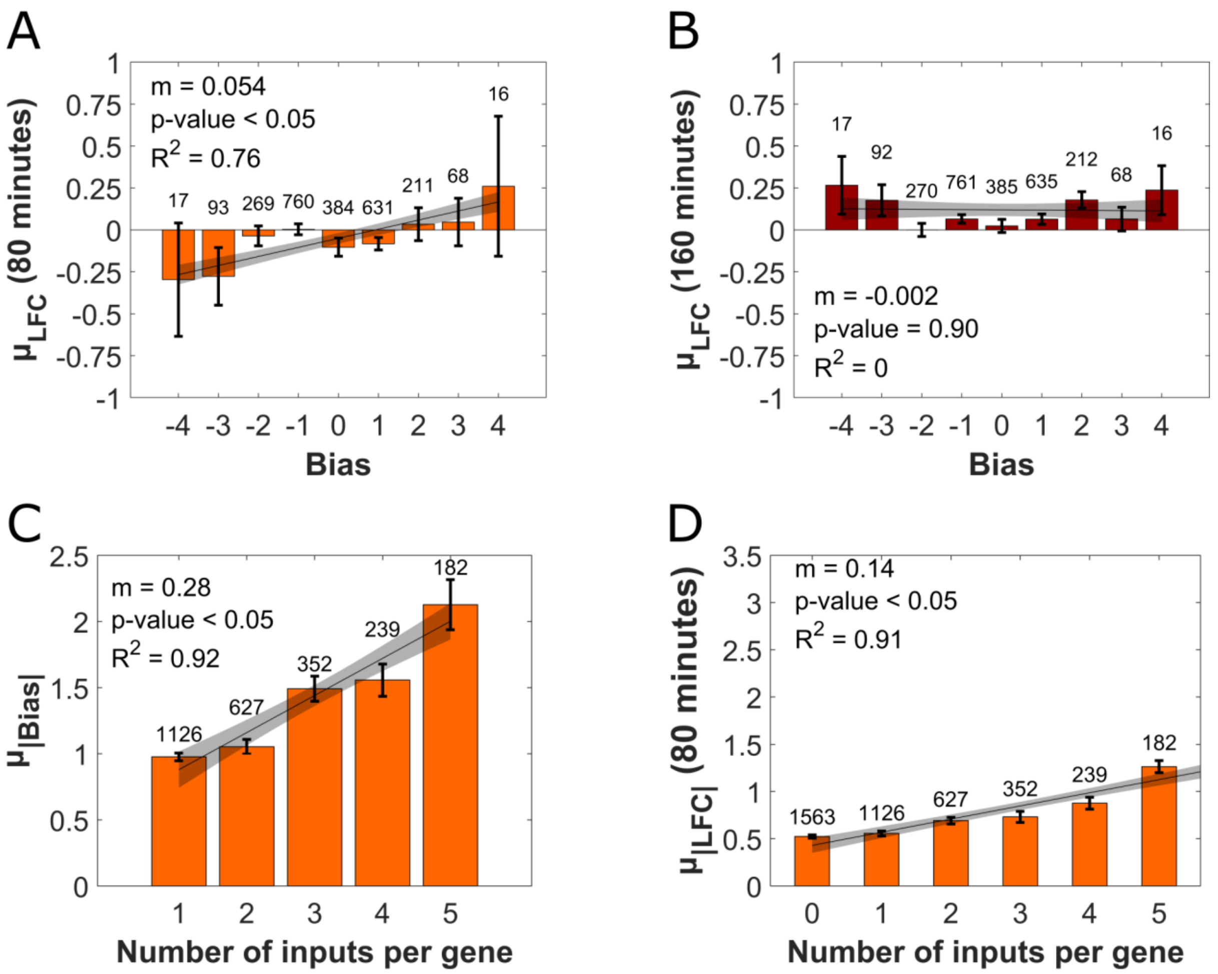


**Figure S2**: TFN and LFCs. **(A)** and **(B)** Average LFCs, $\mu_{LFC}$, of gene cohorts as a function of their mean absolute bias $\mu_{|b|}$, at 80 and 160 minutes, respectively. **(C)** Mean absolute bias, $\mu_{|b|}$, of genes with a given number of input TFs, $K_{TF}$. The error bars are the standard error of the mean (SEM). **(D)** Average absolute LFCs, |LFC|, as a function of $K_{TF}$ at 80 minutes. The error bars are the standard error of the mean. The best fitting line and 68% confidence boundaries (CB, grey) were obtained using FITLM (MATLAB). Also shown are the coefficient of determination $R^2$, inclination of the line *m*, and p-value under the null hypothesis that the lines are horizontal. The hypothesis is not rejected for p value < 0.05. The numbers on top of the bars are the number of genes of the cohort.

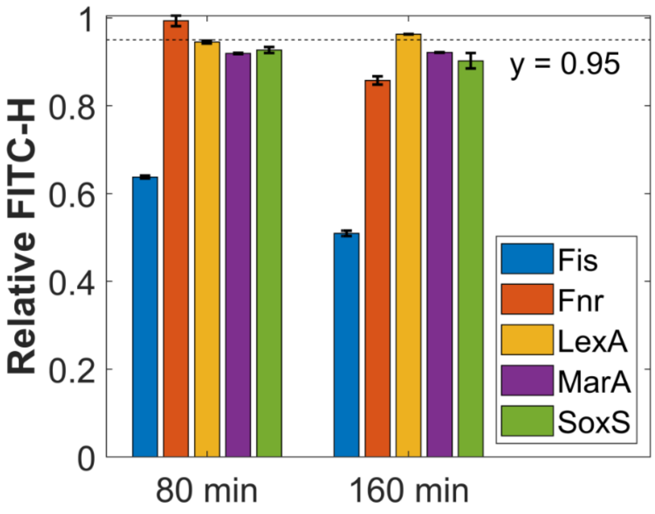


**Figure S3**: Mean single-cell expression levels relative to the control of the GRs Fnr, Fis, LexA, MarA, and SoxS at 80 and 160 minutes after adding rifampicin (measured by flow-cytometry). Error bars are the standard error of the mean (SEM) from 3 biological replicates. The dashed horizontal line at y = 0.95 marks a 5% difference from 1, below which we conclude that it differs significantly from 1 and, thus, that it differs from the control.

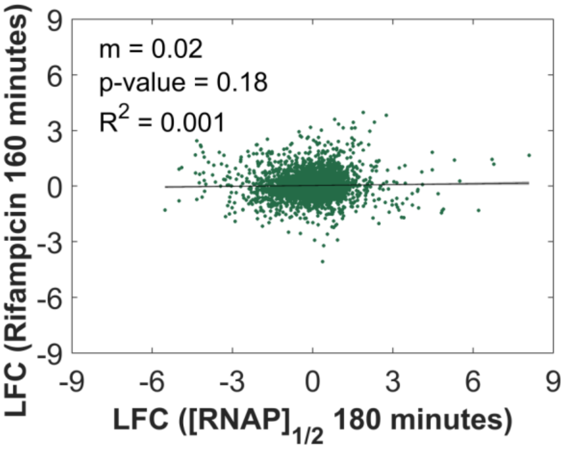


**Figure S4**: Scatter plot of gene response strengths (LFC) to rifampicin at 80 minutes plotted against the same gene's response strength (LFC) to a reduction to half in the RNAP concentration ($[RNAP]_{1/2}$, data from (Almeida2022)). Shown also is the best fitting line and its inclination m, coefficient of determination $R^2$, 95% confidence bounds (small shadow areas), and p-value under the null hypothesis that the line is horizontal (not rejected if < 0.05).

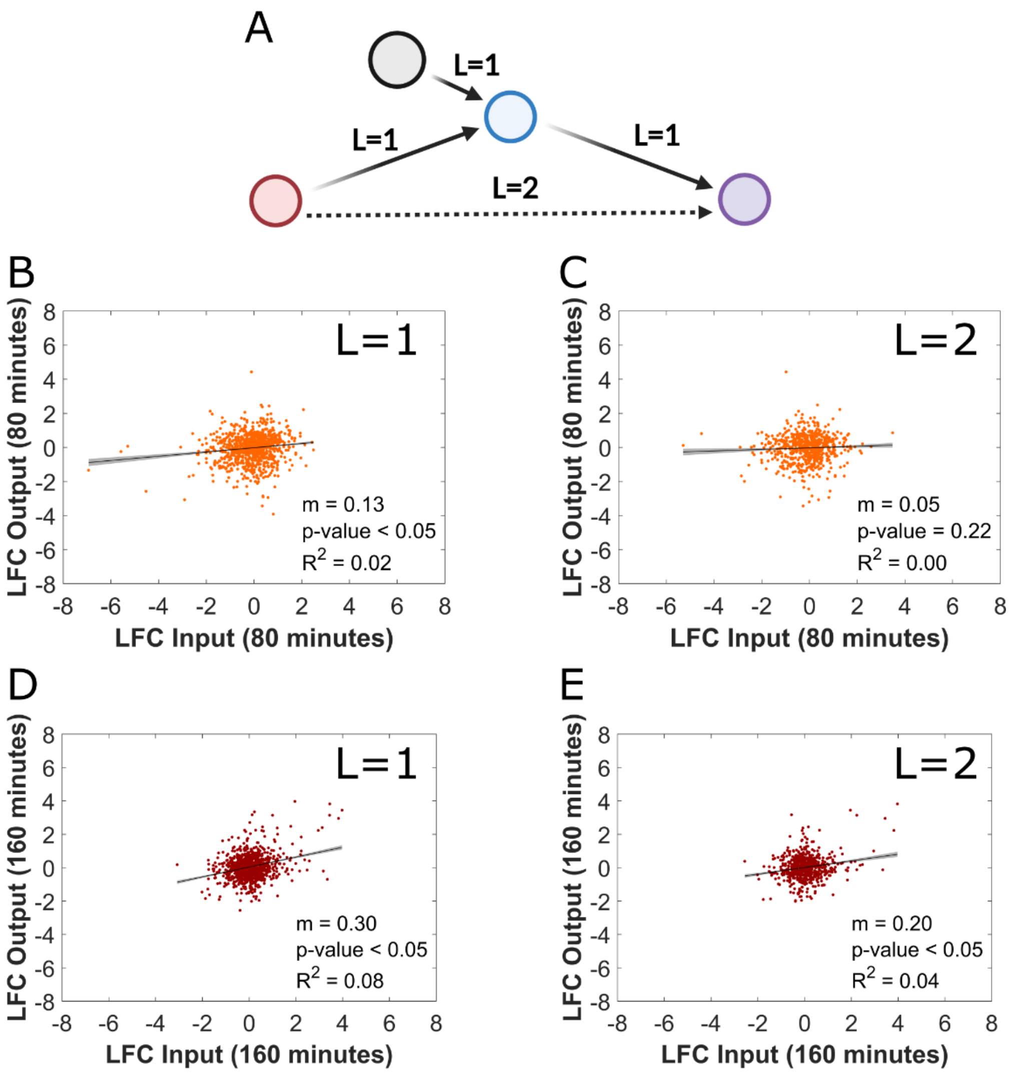


**Figure S5. Input and output TFs response to rifampicin**. **(A)** Illustration of the signal propagation via TFs as a function of the path length (L = 1, 2, …) between genes. The straight lines represent direct links by TFs (L=1), while the dashed line is an indirect link (L=2), made possible by the straight lines. **(B)** and **(C)** are scatter plots of the LFC of TF output genes and TF input genes for a path length L of 1 and 2, respectively, at 80 minutes after adding rifampicin. **(D)** and **(E)** are the same as **(B)** and **(C)** but for 180 minutes after adding rifampicin. Shown also are the best fitting lines and their inclination m, coefficient of determination $R^2$, 95% confidence bounds (small shadow areas), and p-value under the null hypothesis that the line is horizontal (not rejected if < 0.05).

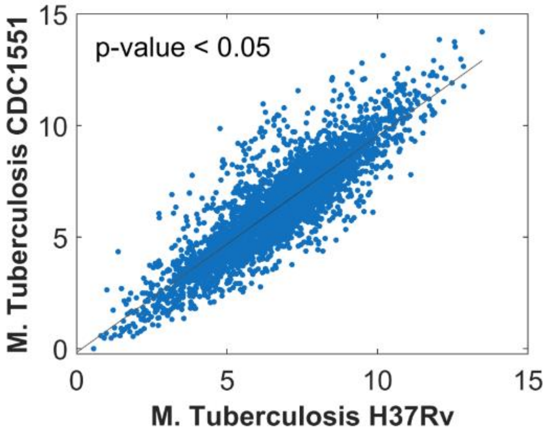


**Figure S6:** Scatter plot of the TPM values of the same genes from two strains of *M. tuberculosis* (H37Rv and CDC1551) 60 minutes under a non-lethal concentration of rifampicin. Shown also is the best fitting line with 95% confidence bounds (very small shadow areas) and the p-value under the null hypothesis that the line is horizontal. The hypothesis is not rejected for $p < 0.05$. The line inclination m equals 0.97 and the coefficient of determination $R^2$ equals 0.77.

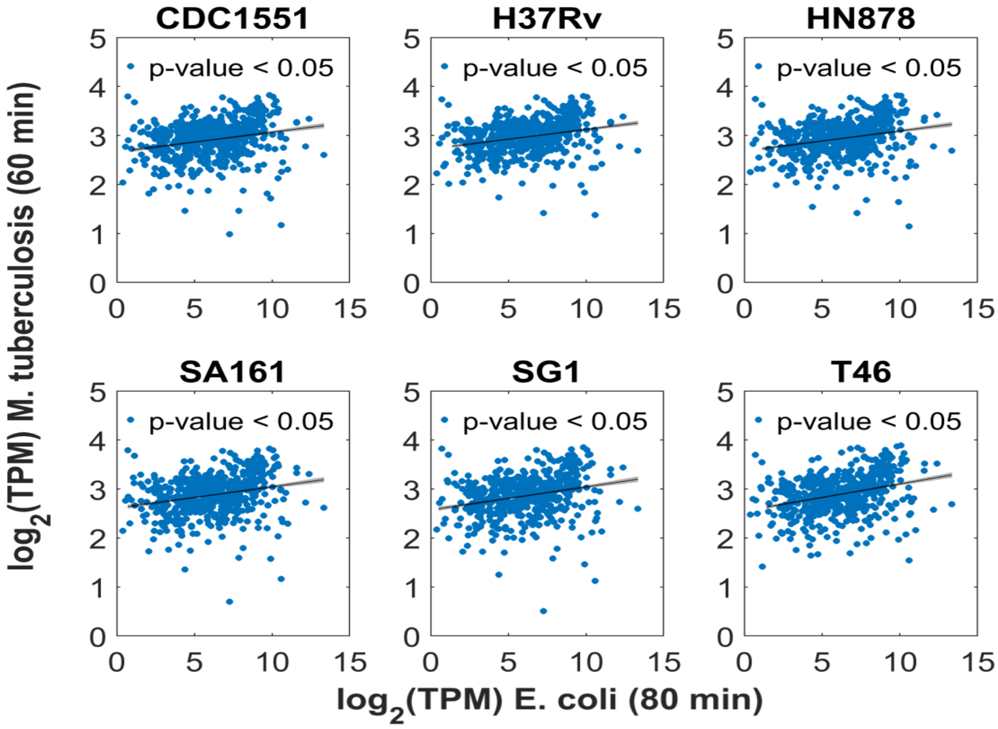


**Figure S7**: Scatter plots of the TPMs of orthologous genes of *E. coli* and various *M. tuberculosis* strains (named on top of each graph) after 80 and 60 minutes under non-lethal concentration of rifampicin, respectively. Shown also are best fitting lines, the 95% confidence bounds (very small shadow areas), and the p-value under the null hypothesis that the line is horizontal. The hypothesis is not rejected for p value < 0.05. Supplementary Table S5 shows the values of m and $R^2$ of each best fit.

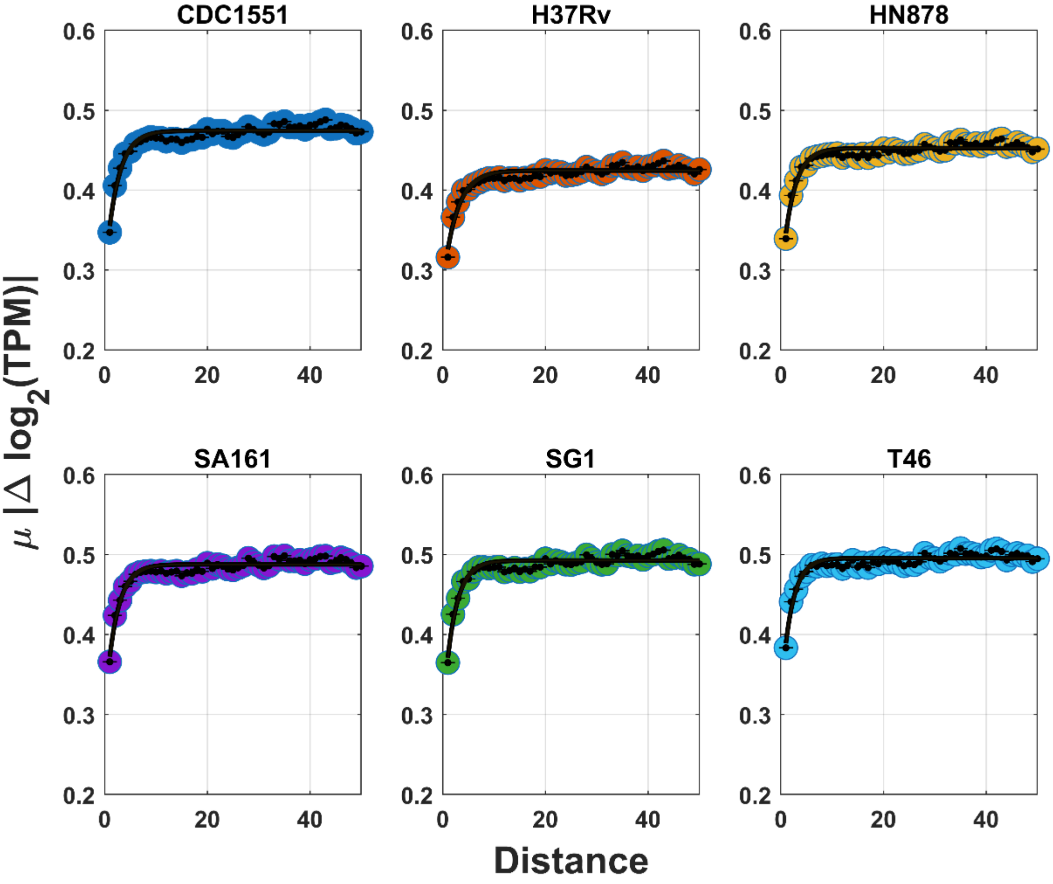


**Figure S8**: Mean absolute differences between the log2(TPM) ($\mu_{|log_2(TPM)|}$) of pairs of genes of *M. tuberculosis* as a function of the distance, D, which equals the number of genes between them in the DNA. Data at 60 minutes under rifampicin. Also shown are best exponential fitting functions: $y = c + a \times e^{-bx}$. The line in each datapoint is an error bar. Supplementary Table S6 shows parameter values and $R^2$ of the fitting functions.

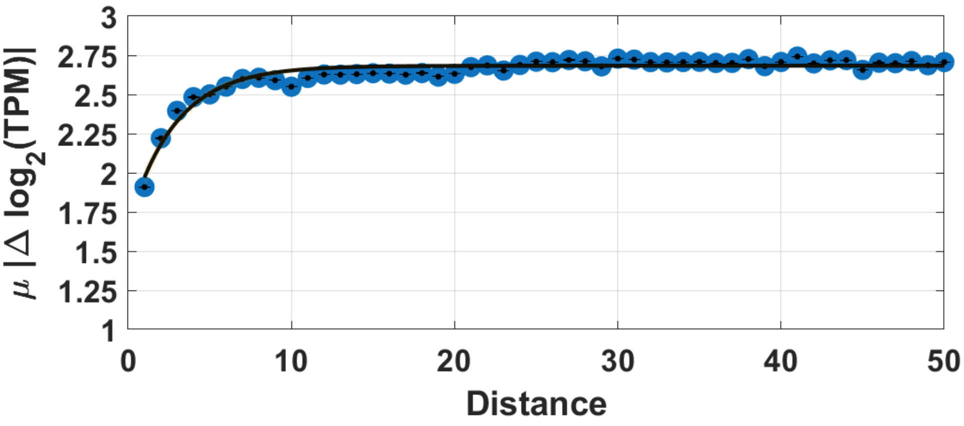

**Figure S9**: Mean absolute differences between the $Log_2$(TPM) ($\mu_{|log_2(TPM)|}$) of all pairs of genes of *E. coli* as a function of the distance, D, which equals the number of genes between them in the DNA. Data from *E. coli*, 80 minutes under rifampicin. Also shown is the best exponential fitting function: $y = 2.7-1.01 \times e^{-0.35.x}$ ($R^2$=0.87 and Adjusted $R^2$=0.87). The black line inside the ball of each datapoint is a (small) error bar.

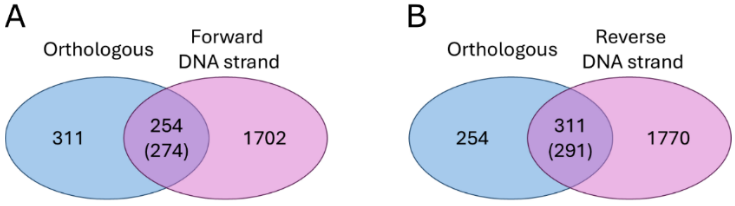


**Figure S10:** Venn diagrams of the number of *E. coli* genes that have orthologous pairs in *M. tuberculosis*, along with the number of genes that are in the **(A)** forward or **(B)** reverse DNA strand. The intersecting areas show the number of genes with both properties. Inside the parentheses are the number of genes expected by random chance to have both properties. From **(A)** the number of genes in the forward DNA strand according to the empirical data is 254, while the number expected by chance is 274, which differs by ~8%.

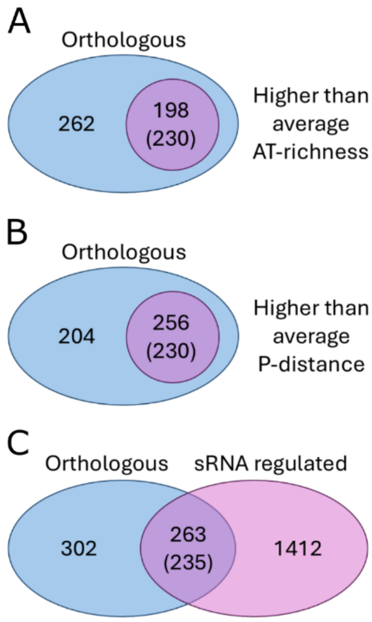


**Figure S11:** Venn diagrams of the number of *E. coli* genes that have orthologous pairs in *M. tuberculosis* as well as other properties. **(A)** Number of orthologous genes that have (198) and that do not have higher than average AT-richness (262). The number of genes with both properties (198) is smaller than expected by random chance. **(B)** Number of orthologous genes that have (256) and that do not have (204) higher than average p-distance. **(C)** Number of orthologous genes that are (263) and that are not regulated by sRNAs (302), along with the number of sRNA regulated genes that are not orthologous (1412). In all cases, inside the parentheses is the number of genes expected to have both properties by random chance.

**Supplementary Tables**

**Table S1:** Differences in RNA levels from the control of genes coding for σ factors as well as of the cohort of genes that they regulate, at 80 minutes after adding rifampicin. μ and STD are the mean and standard deviation of LFC values of σ-regulated genes, respectively. N is the number of those genes. Finally, $p_1$ is the probability that the distribution is the same as the distribution of all non-σ regulated genes and is given by the ttest2 MATLAB function, while $p_2$ is the p-value of the LFC value.

| Regulated genes | μ(LFC) | STD(LFC) | N | $p_1$ | Regulator | LFC | $p_2$ |
|---|---|---|---|---|---|---|---|
| All σ regulated genes | 0.04 | 0.74 | 1980 | - | - | - | - |
| $\sigma^{19}$ regulated genes | -0.18 | 0.45 | 5 | 0.89 | $\sigma^{19}$ | 0.78 | 0.19 |
| $\sigma^{24}$ regulated genes | -0.10 | 0.61 | 114 | 0.94 | $\sigma^{24}$ | -1.59 | <0.05 |
| $\sigma^{28}$ regulated genes | -0.37 | 0.87 | 70 | 0.05 | $\sigma^{28}$ | 0.01 | 0.99 |
| $\sigma^{32}$ regulated genes | -0.09 | 0.54 | 159 | 0.86 | $\sigma^{32}$ | -0.75 | <0.05 |
| $\sigma^{38}$ regulated genes | -0.86 | 1.51 | 317 | <0.05 | $\sigma^{38}$ | -0.31 | <0.05 |
| $\sigma^{54}$ regulated genes | -0.55 | 1.61 | 133 | <0.05 | $\sigma^{54}$ | -0.10 | 0.36 |
| $\sigma^{70}$ regulated genes | -0.02 | 1.08 | 1574 | <0.05 | $\sigma^{70}$ | -0.19 | <0.05 |

**Supplementary Table S2:** Differences in RNA levels from the control of genes coding for σ factors as well as of the cohort of genes that they regulate, at 160 minutes after adding rifampicin. μ and STD are the mean and standard deviation of LFC values of σ-regulated genes, respectively. N is the number of those genes. Finally, $p_1$ is the probability that the distribution is the same as the distribution of all non-σ regulated genes and is given by the ttest2 MATLAB function, while $p_2$ is the p-value of the LFC value.

| Regulated genes | μ(LFC) | STD(LFC) | N | $p_1$ | Regulator | LFC | $p_2$ |
|---|---|---|---|---|---|---|---|
| All σ factor regulated genes | 0.04 | 0.74 | 1989 | - | - | - | - |
| $\sigma^{19}$ regulated genes | 0.24 | 0.18 | 5 | 0.53 | $\sigma^{19}$ | 0.98 | <0.05 |
| $\sigma^{24}$ regulated genes | -0.01 | 0.58 | 116 | 0.55 | $\sigma^{24}$ | -1.03 | <0.05 |
| $\sigma^{28}$ regulated genes | -0.03 | 0.47 | 71 | 0.47 | $\sigma^{28}$ | 0.44 | 0.51 |
| $\sigma^{32}$ regulated genes | -0.41 | 0.64 | 159 | <0.05 | $\sigma^{32}$ | -0.53 | <0.05 |
| $\sigma^{38}$ regulated genes | -0.05 | 0.73 | 316 | <0.05 | $\sigma^{38}$ | -0.25 | 0.23 |
| $\sigma^{54}$ regulated genes | 0.22 | 0.79 | 133 | <0.05 | $\sigma^{54}$ | 0.22 | 0.38 |
| $\sigma^{70}$ regulated genes | 0.06 | 0.77 | 1580 | 0.43 | $\sigma^{70}$ | 0.10 | 0.64 |

**Supplementary Table S3:** Mean μ and standard deviation STD of the LFC of the output genes of GRs, respectively, at 80 minutes. N is the number of genes and $p_1$ is the probability that the distribution is the same as the distribution of all other genes (ttest2 MATLAB function). Also shown are the changes in RNA levels (LFC) of the GRs with rifampicin, at 80 minutes, followed by $p_2$, the p-value of their LFC.

| Global regulator | μ(LFC) | STD (LFC) | N | $p_1$ | GR (LFC) | $p_2$ |
|---|---|---|---|---|---|---|
| crp | 0.97 | 0.99 | 551 | <0.05 | 0.06 | 0.62 |
| fnr | 1.01 | 1.00 | 310 | <0.05 | 0.40 | <0.05 |
| fis | 0.77 | 0.84 | 220 | <0.05 | 1.28 | <0.05 |
| narL | 0.78 | 0.59 | 122 | <0.05 | 0.01 | 0.93 |
| fur | 0.95 | 0.94 | 262 | <0.05 | 0.02 | 0.88 |
| lrp | 0.75 | 0.74 | 348 | <0.05 | 0.03 | 0.81 |
| nsrR | 0.61 | 0.54 | 80 | 0.63 | 0.08 | 0.62 |
| cra | 0.63 | 0.74 | 128 | 0.77 | 0.71 | <0.05 |
| flhD | 0.53 | 0.37 | 83 | 0.10 | 0.92 | <0.05 |

| | | | | | | |
|---|---|---|---|---|---|---|
| cpxR | 0.65 | 0.74 | 115 | 0.99 | 0.41 | <0.05 |
| narP | 0.88 | 0.45 | 50 | <0.05 | 0.29 | 0.05 |
| phoB | 0.83 | 0.69 | 61 | <0.05 | 0.12 | 0.49 |
| lexA | 0.33 | 0.32 | 59 | <0.05 | 0.13 | 0.27 |
| phoP | 0.71 | 0.63 | 54 | 0.50 | 0.30 | <0.05 |
| marA | 0.89 | 1.18 | 56 | <0.05 | 0.09 | 0.74 |
| soxS | 0.59 | 0.52 | 49 | 0.51 | 0.05 | 0.84 |

**Supplementary Table S4:** Mean μ and standard deviation STD of the LFC of the output genes of GRs, respectively, at 160 minutes. N is the number of genes and $p_1$ is the probability that the distribution is the same as the distribution of all other genes (ttest2 MATLAB function). Also shown are the changes in RNA levels (LFC) of the GRs with rifampicin, at 160 minutes, followed by $p_2$, the p-value of their LFC.

| Global regulator | μ (LFC) | STD (LFC) | N | $p_1$ | GR (LFC) | $p_2$ |
|---|---|---|---|---|---|---|
| crp | 0.63 | 0.60 | 550 | <0.05 | 0.37 | 0.11 |
| fnr | 0.61 | 0.56 | 309 | <0.05 | 0.01 | 0.94 |
| fis | 0.73 | 0.60 | 220 | <0.05 | 0.29 | 0.29 |
| narL | 0.51 | 0.45 | 122 | 0.71 | 0.43 | 0.07 |
| fur | 0.61 | 0.61 | 262 | <0.05 | 0.22 | 0.34 |
| lrp | 0.57 | 0.48 | 347 | <0.05 | 0.33 | 0.10 |

| | | | | | | |
|---|---|---|---|---|---|---|
| nsrR | 0.44 | 0.41 | 81 | 0.33 | 0.08 | 0.77 |
| cra | 0.55 | 0.45 | 128 | 0.19 | 0.52 | <0.05 |
| flhD | 0.51 | 0.50 | 83 | 0.82 | 0.53 | 0.28 |
| cpxR | 0.48 | 0.55 | 116 | 0.71 | 0.14 | 0.49 |
| narP | 0.55 | 0.49 | 122 | 0.44 | 0.09 | 0.69 |
| phoB | 0.56 | 0.41 | 62 | 0.34 | 0.22 | 0.36 |
| lexA | 0.37 | 0.36 | 59 | <0.05 | 0.76 | <0.05 |
| phoP | 0.59 | 0.52 | 54 | 0.15 | 0.38 | 0.07 |
| marA | 0.65 | 0.58 | 54 | <0.05 | 0.72 | <0.05 |
| soxS | 0.45 | 0.37 | 50 | 0.48 | 0.11 | 0.77 |

**Supplementary Table S5:** Parameter values m, p-value, and $R^2$ of the fitting lines to the correlation plots between the TPMs of orthologous genes of *E. coli* and *M. tuberculosis* after 80 and 60 minutes under non-lethal concentration of rifampicin, respectively.

| **Strain** | **m** | **p value** | **$R^2$** |
|---|---|---|---|
| CDC1551 | 0.04 | < 0.05 | 0.04 |
| H37Rv | 0.04 | < 0.05 | 0.06 |
| HN878 | 0.04 | < 0.05 | 0.05 |
| SA161 | 0.04 | < 0.05 | 0.05 |
| SG1 | 0.05 | < 0.05 | 0.05 |
| T46 | 0.05 | < 0.05 | 0.08 |

**Supplementary Table S6:** Parameter values of the exponential fitting functions in Supplementary Figure S8: $y = c + a \times e^{-bx}$, where a is the amplitude (initial offset from the baseline), b is the growth rate, and c is the saturation as x goes to infinity. The fitting was performed using a MATLAB fit function with a custom-defined model (using 'fit type'). The parameters a, b, and c were estimated by nonlinear least squares regression. Each model was evaluated in 100 iterations, using different starting points to ensure convergence to a global or near-global minimum. The quality of each fit was assessed by the coefficient of determination ($R^2$) and the adjusted $R^2$, which account for both the number of fitted parameters and the sample size. From 100 fits, the model with the highest $R^2$ was selected as the best fitting curve. In all cases, the model with the highest $R^2$ also had the highest adjusted $R^2$, supporting a consistently strong fit to the data.

| Strain | $R^2$ | Adjusted $R^2$ | a | b | c |
|---|---|---|---|---|---|

| | | | | | |
|---|---|---|---|---|---|
| CDC1551 | 0.88 | 0.88 | -0.18 | 0.42 | 0.47 |
| H37Rv | 0.87 | 0.87 | -0.14 | 0.38 | 0.42 |
| HN878 | 0.87 | 0.87 | -0.16 | 0.45 | 0.45 |
| SA161 | 0.88 | 0.88 | -0.18 | 0.45 | 0.49 |
| SG1 | 0.88 | 0.88 | -0.19 | 0.46 | 0.49 |
| T46 | 0.86 | 0.86 | -0.18 | 0.50 | 0.50 |
